# A versatile toolkit for CRISPR-Cas13-based RNA manipulation in *Drosophila*

**DOI:** 10.1101/2020.09.25.314047

**Authors:** Nhan Huynh, Noah Depner, Raegan Larson, Kirst King-Jones

**Affiliations:** Department of Biological Sciences University of Alberta, G-504 Biological Sciences Bldg., Edmonton, Alberta T6G 2E9, Canada

**Keywords:** CRISPR, Cas13, CasRX, *Drosophila*, RNA manipulation, crRNA design

## Abstract

Advances in CRISPR technology have immensely improved our ability to manipulate nucleic acids, and the recent discovery of the RNA-targeting endonuclease Cas13 adds even further functionality. Here, we show that Cas13 works efficiently in *Drosophila*, both *ex vivo* and *in vivo*. We tested 44 different Cas13 variants to identify enzymes with the best overall performance and showed that Cas13 could target endogenous *Drosophila* transcripts *in vivo* with high efficiency and specificity. We also developed Cas13 applications to edit mRNAs and target mitochondrial transcripts. Our vector collection represents a versatile tool collection to manipulate gene expression at the post-transcriptional level.

## Background

Most bacterial and archaeal genomes harbor <u>C</u>lustered <u>R</u>egularly Interspaced <u>S</u>hort <u>P</u>alindromic <u>R</u>epeats (CRISPR) and encode <u>C</u>RISPR-<u>a</u>ssociated protein<u>s</u> (Cas) as a defense system against bacteriophages and other invading nucleic acids [1–3]. The immune response of all CRISPR/Cas systems characterized to date includes three steps: i) adaptation and spacer acquisition, where a piece of the invading genome is incorporated into the CRISPR array; ii) the expression of mature CRISPR RNAs (gRNAs) from the processed CRISPR array and iii) interference, where Cas enzymes are guided by the gRNAs to the corresponding region of the invading genome for cleavage and degradation [4, 5]. The CRISPR/Cas class II systems use a single, multidomain Cas effector protein [6]. Because of its simplicity, the single multidomain effector found in class II organisms is used in current CRISPR methods. Class II type II CRISPR Cas9 was one of the first Cas proteins studied in detail, which led to its widespread use for genomic engineering (Figure 1A) [6–11]. Currently, CRISPR/Cas9 approaches allow scientists to precisely alter gene function via i) classic CRISPR to introduce short INDELs, ii) HR-based CRISPR for homology-based gene replacements or deletions, iii) somatic CRISPR for conditional gene disruption, iv) CRISPRi, (i = interference) to interfere with gene transcription, and v) CRISPRa (a = activation) to upregulate gene activity. Studies have shown that it is possible to conditionally target genes of interest by exerting spatial and temporal control over Cas9 expression or using ligand-activated Cas9 variants [8,10,12,13]. The rapid advances in CRISPR technologies have made it a popular choice over earlier nuclease-based gene editing approaches like meganucleases (MNs) [14, 15], zinc finger nucleases (ZFNs) [16–18], and transcription activator-like effector nucleases (TALENs) [19, 20].

**Figure 1:**
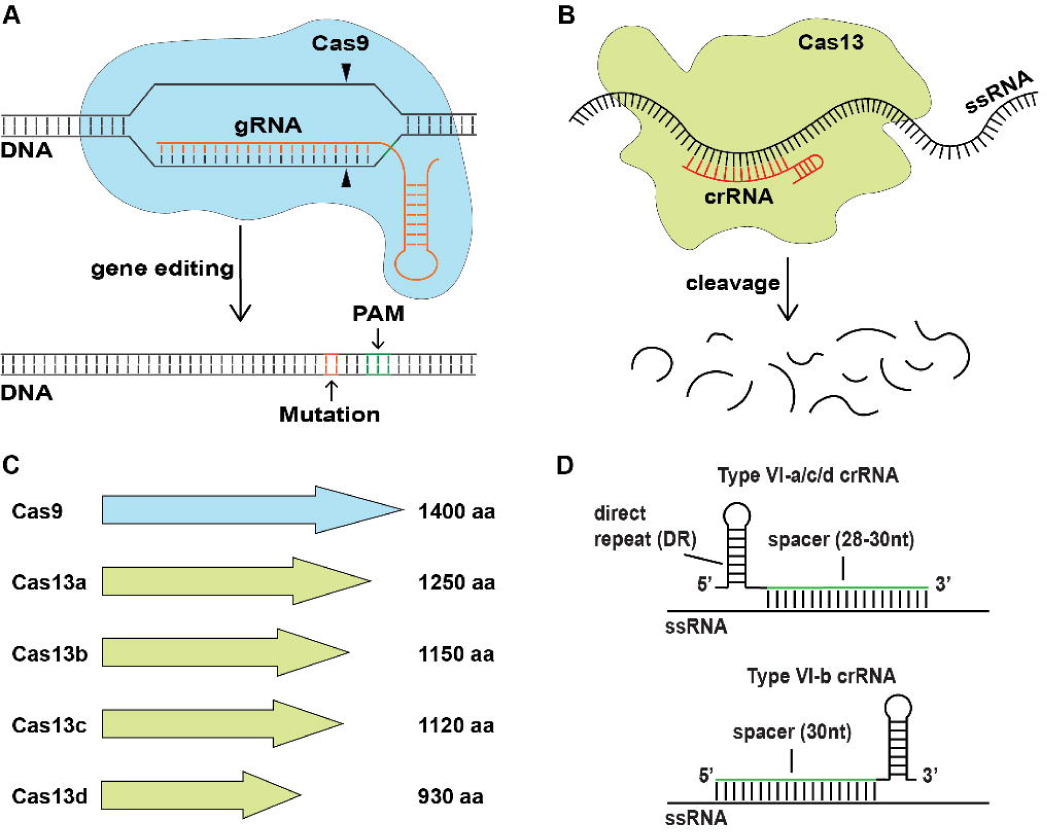
Functional overview of CRISPR/Cas9 and CRISPR/Cas13 systems. **(A)** Schematic of Cas9 mechanism in genome editing. This system requires the recruitment of CRISPR-associated protein Cas9 (blue) to the target site recognized by the guide RNA (gRNA: orange). Target site cleavage by Cas9 is ensured by the presence of the protospacer adjacent motif (PAM) (green), a sequence that immediately follows the target site. The PAM will determine the Cas9 cleavage site, which lies about three nucleotides upstream of the PAM. **(B)** Schematic of the Cas13 RNA cleavage mechanism. This system requires the pre-assembly of Cas13 (green) with the CRISPR RNA (crRNA: red) to recognize target RNAs. Upon RNA-binding, Cas13 will undergo a conformational change and induce the catalytic activity of its nuclease domains, resulting in the cleavage of target transcripts. **(C)** Comparisons of Cas9 size with different Cas13 subtypes (a-d). Polypeptide sizes are indicated as the number of amino acids. **(D)** Relative structural representation of different Cas13 subtype-compatible crRNAs. All four subtype crRNAs carry a direct repeat to facilitate the binding with their corresponding Cas13 enzyme, as well as a spacer sequence specific for the target transcript. Cas13b-compatible crRNAs carry a direct repeat at the 3’-end while compatible crRNAs for Cas13a, c, and d carry the direct repeat at the 5’-end.

The recent introduction of the class II type VI CRISPR/Cas13 system further expands the existing technology in significant ways. Like Cas9, Cas13 uses a guide RNA (CRISPR-RNA, aka crRNA) to identify its substrate, which is RNA rather than DNA (Figure 1B). Cas13 enzymes have two distinct catalytic activities: i) an RNAse activity that is mediated by two <u>h</u>igher <u>e</u>ukaryotic and <u>p</u>rokaryotic <u>n</u>ucleotide (HEPN) binding domains and ii) a gRNA maturation activity, possibly a combination of activities located in the HEPN2 and Helical-1 domains [21, 22]. There are currently four subtypes identified in the Cas13 family, including Cas13a (aka C2c2), Cas13b, Cas13c, and Cas13d. All Cas13 family members are smaller than Cas9, with Cas13d being the smallest protein. The small size of Cas13 proteins makes them suitable for molecular genetics (Figure 1C). All Cas13 enzymes require a 60-66 nucleotide long crRNA to ensure target specificity [2,3,23]. Similar to the gRNA in the CRISPR/Cas9 system, the crRNA used by Cas13 forms a short hairpin structure next to a short spacer sequence (28-30 nucleotides) that is specific to the target transcript (Figure 1D). Since CRISPR/Cas13 mediates RNA degradation, it holds the promise to replace or complement RNA interference (RNAi) approaches or other systems that interfere with transcript levels, such as CRISPRi. Despite being a powerful tool, RNAi often suffers from low efficiencies or off-target effects, whereas Cas9-based CRISPRi requires a <u>p</u>rotospacer <u>a</u>djacent <u>m</u>otif (PAM), thus limiting the flexibility by which target sequences can be selected [24–28]. It is desirable to examine whether CRISPR/Cas13 can offer better specificity and efficiency than these other interference techniques.

*Drosophila melanogaster* is a versatile genetic model organism that is used to study a wide variety of biological processes. Traditional techniques to analyze gene function in *Drosophila* include the generation of mutations via chemical mutagens and transposable P-elements, or the use of transgenes to trigger RNAi and to express cDNAs for gain-of-function studies via the Gal4/UAS system [29–33]. Like other model organisms, the CRISPR/Cas9 endonucleases have been quickly adopted by *Drosophila* researchers [10,24,25,34–39]. CRISPR-based techniques are remarkably precise and, therefore, ideal for replacing, validating, and complementing traditional approaches, in particular procedures relying on the expression of RNAi or cDNA transgenes [40, 41]. Also, the large worldwide collection of gRNAs stocks has ensured the quick adaptation of CRISPR/Cas9 into mainstream *Drosophila* research [6,42,43]. Given the potential of CRISPR/Cas13-based methods to replace current techniques, we explored its feasibility and reliability in *Drosophila*.

Our lab studies signaling pathways that control ecdysone and heme biosynthesis in the larval prothoracic gland (PG), which is part of a larger structure called the ring gland. The PG is a popular model for investigating fundamental aspects of insect endocrinology and allows for the study of external cues that control the timing of ecdysone pulses [44]. Recently, we carried out a genome-wide PG-specific RNAi screen that identified 1,906 genes with critical roles in larval development [45]. In follow-up experiments, however, we often were unable to validate the RNAi-induced phenotypes by independent RNAi lines, either because no such lines existed or because other RNAi lines did not replicate the phenotype. This prompted us to develop CRISPR-based methods that could validate the RNAi results by an unrelated methodology. We previously generated two CRISPR/Cas9 toolkit collections and could use them to validate some RNAi phenotypes. However, specific issues still exist, including inconsistent gRNA efficiency and early lethality. We sought to investigate the possibility of adapting the CRISPR/Cas13 system for interference and other potential applications of this system in *Drosophila melanogaster*.

We generated and evaluated the catalytic activity of *Drosophila* codon-optimized Cas13 (a-d) variants in a cell line derived from Sg4 embryonic cells. We refer to these Cas13 variants as CasFA[n], CasFB[n], CasFC[n] and CasFX[n], respectively (F = Fruit fly, A-C indicates the Cas13 subfamily, CasFX is the fly version of CasRX, and [n] indicates variant number)(Figure 2A-D). “CasRX” was coined by Konermann et al. for the Cas13d ortholog isolated from *Ruminococcus flavefaciens* XPD3002 to distinguish it from other Cas13d variants [46]. Since we generated fly-optimized versions of CasRX, we refer to these versions as CasFX. Once we had identified a fly-optimized Cas13 variant, we used this variant to adapt existing Cas13 mammalian cell culture applications for *Drosophila* cells, such as transcript tracking and RNA modification [23,47–52]. These *ex vivo* procedures formed the basis for generating a collection of transgenic CRISPR/Cas13 tools designed for *in vivo* RNA targeting. In particular, we generated four Cas13 transgenic lines, namely two that either ubiquitously express CasFB or CasFX, and two that express either CasFB or CasFX under UAS control. The UAS lines allow tissue-specific expression of CasFX and CasFB by crossing them to Gal4-expressing flies. As proof-of-principle that these Cas13 transgenes work effectively *in vivo*, we generated seven crRNA transgenes to target three genes we are studying in our lab.

**Figure 2:**
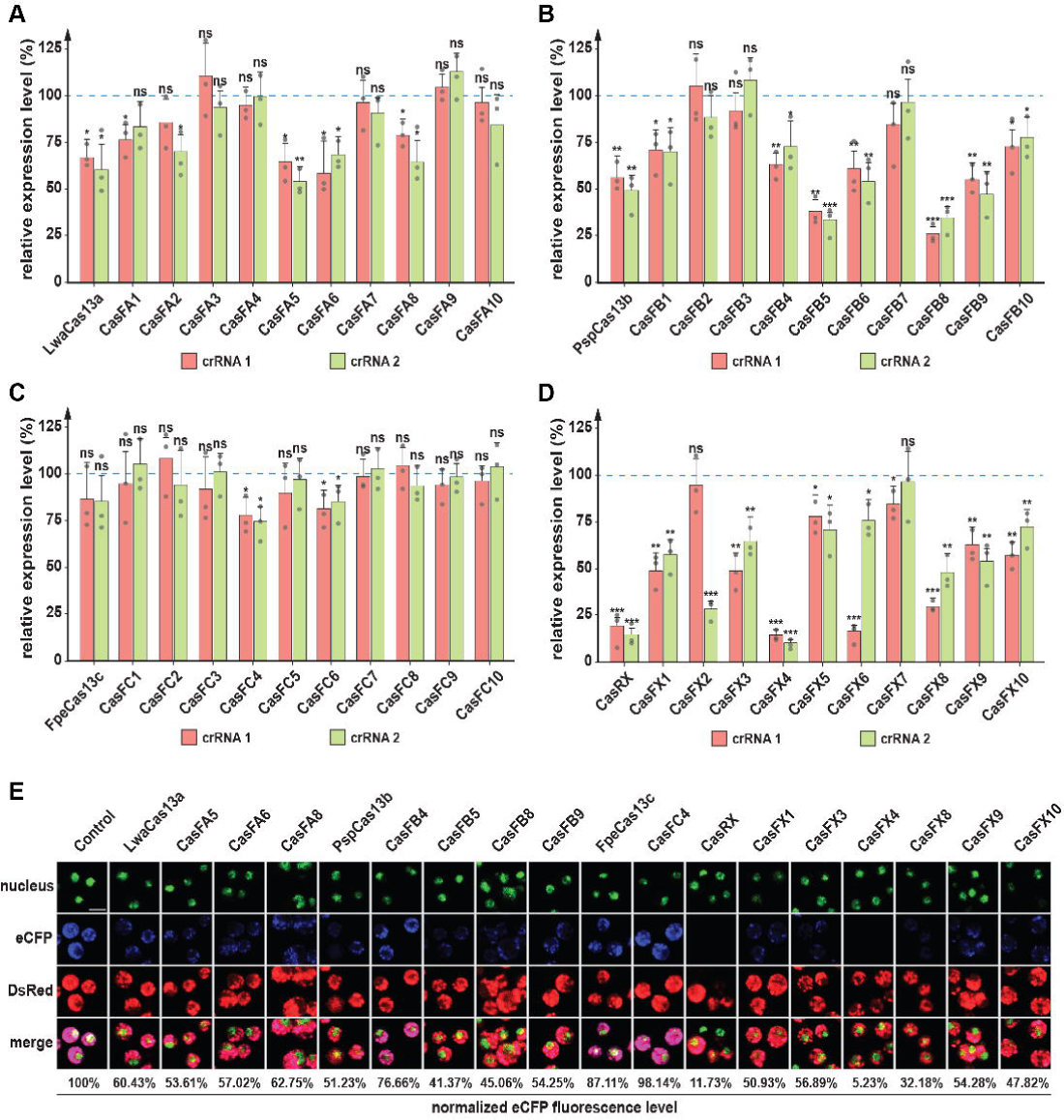
Efficiency evaluation of *Drosophila* codon-optimized Cas13 variants. **(A-D)** qPCR analysis showing eCFP transcript levels in Sg4 cells as a function of the different Cas13 variants that were expressed in these cells (a-d, respectively). Shown are relative fold changes of eCFP transcript being targeted by two independent crRNAs, crRNA 1 (red) and crRNA 2 (green). Data were normalized to eCFP expression levels when using a blank crRNA as a control (blue dotted line = 1). * = p-value < 0.05, ** = p-value < 0.01, *** = p-value < 0.001, ns= not significant, p-values based on Student’s t-test, error bars represent 95% confidence intervals. **(E)** Fluorescence changes of eCFP across samples targeted by the Cas13/crRNA 2 complex. Fluorescence levels were measured using ImageJ and normalized to signals obtained with a blank crRNA (control). Nuclei were stained with Nuclear Green LCS1 (ab138904). eCFP and DsRed fluorescence were measured using their native fluorescence properties (no antibody staining). Scale bar = 50 μm.

## Results

### Generation and characterization of *Drosophila-*optimized Cas13 enzymes

We generated ten Cas13 variants for each of the four Cas13 family members (a-d) by optimizing different codon subsets for codon usage in *Drosophila*. Specifically, we made ten constructs based on the *Leptotrichia wadei Cas13a* gene (*LwaCas13a*), ten variants based on the *Prevotella* sp. P5-125 *Cas13b* gene (*PspCas13b*), ten versions based on the *Fusobacterium perfoetens Cas13c* gene (*FpeCas13c*) and ten forms of the *Ruminococcus flavefaciens* XPD3002 Cas13d gene (aka CasRX) (Table S1). We chose these Cas13 orthologs for the following reasons: i) based on studies in mammalian and plant cells, LwaCas13a, PspCas13b and CasRX showed improved and robust catalytic efficiency when compared to other Cas13 orthologs [23,46,50,53], ii) unlike some Cas13 orthologs, the Cas13 genes we chose for our studies do not require a specific <u>p</u>rotospacer flanking <u>s</u>equence (PFS) for efficient target RNA identification [23,46,50,53]. In the case of PspCas13b, the original study, which was performed in *Escherichia coli,* showed that the PFS is necessary for RNA cleavage activity. However, when the same enzyme was tested in mammalian cells and plants, the PFS was no longer required [48,49,52]. Finally, iii) we also selected Cas13c, since only a few studies have examined this Cas13 subtype [23].

To evaluate the RNA degradation efficiency of these fruit fly-optimized Cas13 enzymes, we needed to establish a stable reporter gene cell line. For this, we used the PhiC31 integrase system to generate a dual-reporter transgene in the *Drosophila* embryo cell line Sg4-PP-27F [54] that simultaneously expressed eCFP (enhanced Cyan Fluorescent Protein) and DsRed (*Discosoma* Red fluorescent protein) (Figures S1A, C). Sg4 is one of four embryonic cell lines isolated from the original Schneider’s line 2 (S2) and differs from the popular S2 cells in a range of transcriptional properties [55]. Importantly, Sg4-PP-27F cells were modified from the original Sg4 cells by adding a PhiC31 docking site to the second chromosome [54]. The inserted eCFP and DsRed transgenes are each controlled by the ubiquitous *actin 5C* promoter (*act5C*). To ensure this transgene’s stability, we added a *Neo^R^* gene cassette, which encodes aminoglycoside kinase, and ensures cell survival in the presence of G418 antibiotics [56]. We refer to this new transgenic cell line as Sg4_CD (C = eCFP, D = DsRed), and our subsequent cell culture experiments were based on this line. To transform the Sg4_CD cell line with appropriate vectors, we generated plasmids that harbored a single copy of a given Cas13 variant and a single crRNA (the vector allows for adding multiple crRNAs). These constructs, here referred to as pC13cr01, allowed us to simultaneously express Cas13 as well as its crRNA in transfected cells (Figure S1D, Table S2). To ensure stable transfection, we also included the *PURO* gene in the pC13cr01 vector. The *PURO* gene encodes the puromycin N-acetyltransferase, which allows cells to survive in media supplemented with puromycin [57, 58] (Figure S1D, Table S2). Thus, the presence of two resistance markers allowed for dual selection during the transfection experiments. Besides testing the *Drosophila*-optimized Cas13 variants, we also examined the efficiency of the original Cas13 orthologs in the Sg4_CD cell line (Tables S1, S2).

We measured the efficiency of the Cas13 variants by targeting one of the two reporter gene mRNAs and quantifying mRNA levels via qPCR. To accomplish this, for each Cas13 variant, we used two independent single crRNAs targeting eCFP mRNA (crRNA1 and crRNA2, Figure 2), while the DsRed mRNA was not targeted and served as a control (Figure S2, Table S4). To ensure that any observed differences derived only from the catalytic activity of the Cas13/crRNA complex, and not from either Cas13 or the crRNA itself, we also tested the eCFP expression level in the presence of a non-targeting (NT) Cas13/crRNA complex. In our hands, the different Cas13 variants showed a wide range of RNA-targeting efficiency, with some of the variants failing to trigger RNA degradation. The original Cas13a, (aka LwaCas13a) showed roughly 35-40% eCFP knock-down efficiency, while the best-performing *Drosophila* variant, CasFA5, was only slightly better and exhibited 47% efficiency (Figure 2A). For the Cas13b (aka PspCas13b) variants, we measured 45-51% efficiency for the original Cas13b enzyme, while the best-performing *Drosophila* variants were CasFB5 and CasFB8, both of which were 65-70% efficient (Figure 2B). The Cas13c group was the least efficient in knocking down eCFP, with the best line, CasFC4, only accomplishing a 25% knock-down (Figure 2C). In contrast, the Cas13d group performed best, displaying 82% efficiency for the original Cas13d (CasRX) enzyme, whereas the CasFX4 variant was even better and reached a 90% knock-down (Figure 2D).

To validate these qPCR data, we quantified the protein levels of eCFP and DsRed based on their fluorescence and Western blotting. We selected the best-performing enzyme variants from all four groups, namely three CasFA variants, four CasFB versions, one CasFC enzyme, and six CasFX forms. We then assessed the efficiency of the eCFP knock-down via immunofluorescence (Figure 2E) and Western blotting (Figure S3A-D). Both approaches showed comparable results and confirmed that CasFX4 was the overall most efficient Cas13 enzyme of the entire cohort, showing ∼90% and ∼95% efficiency on the mRNA and protein levels, respectively.

Next, we sought to investigate whether the subcellular localization of Cas13 would affect the enzyme’s catalytic activity. Since mRNAs mature in the nucleus but are translated in the cytoplasm, we wondered if Cas13 performance could be improved by identifying which cellular compartment is optimal for Cas13 activity. To test this, we selected the original Cas13 variants and their corresponding best-performing *Drosophila* counterparts (CasFA5, CasFB8, CasFC4, and CasFX4), and fused them either with a <u>n</u>uclear localization <u>s</u>ignal (NLS) or a <u>n</u>uclear <u>e</u>xport <u>s</u>ignal (NES) (Figure S3E). These constructs were based on similar designs from other studies and our approaches (Figure S3F) [8,10,12,35,39,59–62]. Then, as described above, we again examined how efficiently eCFP was knocked down. Overall, we observed similar efficiencies when the same Cas13 variant was tested in the nucleus or cytoplasm, indicating that the catalytic activity of these Cas13 variants was independent of the subcellular localization (Figure S3G). For LwaCas13a, PspCas13b and CasRX, this result is consistent with a previous study in plants [52]. Since we found no significant differences, we decided to use Cas13 variants without any localization signal for experiments that followed.

Together, these data suggested that the Cas13 variants retain their RNA-cleaving activity in *Drosophila* Sg4_CD cells, but efficiencies varied considerably. Among the *Drosophila* codon-optimized Cas13 enzymes we generated, we noticed consistent and robust efficiency of two CasFB versions (namely CasFB5 and CasFB8) and the overall best Cas13 variant, CasFX4.

### Evaluating the collateral activity of *Drosophila*-optimized Cas13 variants

Studies in *Escherichia coli* showed that once the Cas13/crRNA complex is bound to its target RNA, the HEPN-nuclease domains become active and are capable of cleaving not just the intended target, but also RNA molecules that are in the vicinity of the Cas13/RNA complex, resulting in the non-specific RNA degradation referred to as “collateral activity” (Figure S4A) [21–23,63]. Subsequent studies reported that the collateral activity of Cas13 varied from system to system. While non-specific RNA degradation was detected in human U87 glioblastoma cells [63], no collateral activity was detected in human embryonic kidney 293FT cells or in the plant *Nicotiana benthamiana* [23,49,50]. To test for collateral activity in our hands, we examined the best-performing Cas13 variants using the same transgenic cell line Sg4_CD. Specifically, we co-expressed eCFP, DsRed, and Neo^R^ independently, each with an *act5C* promoter. Since eCFP, DsRed, and aminoglycoside kinase (encoded by *Neo^R^* gene) are foreign genetic components, we reasoned that manipulating their expression via Cas13 would not have a significant impact on the physiology of SG4_CD cells. The idea was to target eCFP with specific crRNAs in the presence of Cas13 and monitor the expression of DsRed as a readout for collateral activity. Both eCFP and DsRed were presumed to be highly expressed in a coordinate fashion, since the act5C promoter controlled each transgene. As such, if the interference activity of Cas13 was not specific to eCFP, we expected to detect differences in DsRed expression via qPCR. Using this approach, our data showed that the selected Cas13/crRNA complexes only affected target-eCFP expression, while DsRed expression appeared unperturbed (Figure S4B). These data suggest that the tested Cas13 enzymes did not have any detectable collateral activity, at least not in the *Drosophila* Sg4_CD cell line.

### Testing the fidelity of *Drosophila* Cas13 variants

Our efforts identified several Cas13 versions that efficiently degraded target RNAs in *Drosophila* cells while exhibiting no detectable collateral activity. Next, we wanted to assess how mismatches between crRNAs and their cognate target RNA would affect RNA degradation as a means to define Cas13 fidelity. In particular, we were curious as to whether Cas13 would display higher fidelity – and as such, lower off-target rates – than RNA interference (RNAi), which is widely used in a variety of research models, ranging from cell culture to whole organisms [64–66]. While RNAi is an attractive and powerful tool, its usability is often hampered by its off-target activity, which can make it challenging to interpret phenotypes, and validation strategies involving codon-modified genes/cDNAs are cumbersome and harbor pitfalls [26,67,68]. Other validation strategies include non-overlapping RNAi constructs targeting distinct regions on the mRNA, classic mutants, or conditional CRISPR/Cas9 approaches. To test the propensity of our Cas13 enzymes to degrade off-target RNAs due to small sequence differences, we selected the six top-performing variants for which we had not detected any collateral activity (CasFA5, CasFB5, CasFB8, CasFC4, CasFX4, and CasFX8). Specifically, we generated mismatches in the crRNA-2 spacer sequence and measured the ability to degrade its target RNA, eCFP. To indicate the mismatch location, we referenced the position of the altered nucleotide relative to the stem loop-forming direct repeat of the crRNA. The nucleotide at position 1 represents the one closest to the DR, and the highest number corresponds to the nucleotide farthest away from the DR.

Among all variants that we tested, all had a central region that appeared to be intolerant to single mismatches. The CasFA5, CasFB5, and CasFB8 variants showed some tolerance to single mismatches outside the core region, namely nucleotides #1-3 at the 5’-end and nucleotides #28 and higher at the 3’-end. In contrast, the core region showed no tolerance to mismatches (Figures 3A, B, C, D). Remarkably, CasFC4, CasFX4 and CasFX8 variants showed no tolerance for mismatches throughout the entire range, including the extreme 5’ and 3’ ends. To examine this further, we tested the outermost nucleotides for both CasFX variants (position #1 and #30). Even single mismatches at either end of the spacer region abrogated interference activity, indicating that these two variants are highly specific and have the lowest off-target potential (Figures 3E, F). Since four of the variants had some tolerance towards a single mismatch, we further examined mismatch tolerance by introducing more than one mutation per crRNA. Specifically, we generated constructs encoding two, three, or four mismatches in the eCFP-crRNA. In all tested conditions, we included at least one mismatch from the extreme 5’ or 3’ end of the spacer. In our hands, none of the *Drosophila* Cas13 variants exhibited tolerance to crRNAs with mismatches of more than one nucleotide (Figures S4 C-H). These data are in agreement with other studies using similar approaches [51,69,70]. Taken together, this suggests that the *Drosophila* Cas13 variants tested here are highly specific and display no tolerance to a single mismatch in the core region of the spacer, and none of the enzymes were functional with two mismatches in the crRNA. The CasFA, CasFB5 and CasFB8 variants did tolerate a single mismatch located at either end outside the core region. In contrast, the CasFC4, CasFX4 and CasFX8 variants appeared to require a perfect match of the entire spacer region to mediate interference. We conclude that the CasFX4 and CasFX8 variants will likely have the lowest off-target rate while retaining optimal RNA-targeting efficiency among the Cas13 enzymes tested here.

**Figure 3:**
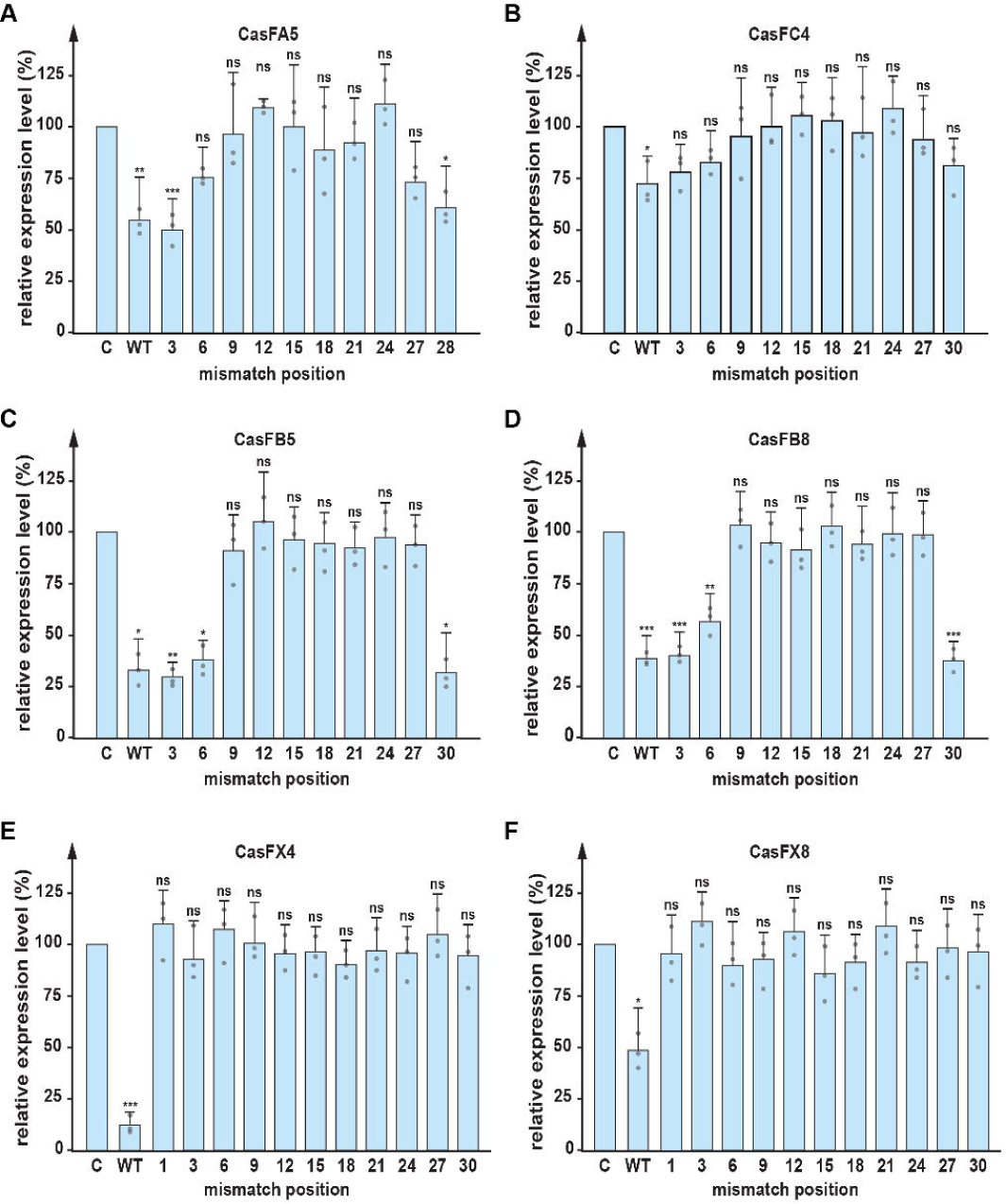
Specificity evaluation of *Drosophila* codon-optimized Cas13 variants in Sg4 cells. **(A-F)** Relative expression levels of eCFP when using different Cas13 variants and crRNAs that carry a range of single mismatches along the eCFP crRNA-2. Data were normalized to samples treated with a blank crRNA (control = C). eCFP expression levels in Cas13/ wild-type (WT) crRNA samples were also included as a reference. * = p-value < 0.05, ** = p-value < 0.01, *** = p-value < 0.001, ns= not significant, p-values based on Dunnett’s *post-hoc* test, error bars represent 95% confidence intervals.

### Nuclease-dead CasFX for applications involving transcript detection

The CRISPR/Cas9 system has been modified to allow for non-nuclease activities, such as for transcription interference (CRISPRi) as well as transcriptional activation (CRISPRa) [8,10,12,39]. Similarly, the Cas13 system can also be adapted for other purposes and may be more suitable for certain applications than CRISPR/Cas9-based methods. For instance, the ability to target RNA instead of DNA has the advantage that it is reversible. Also, Cas13 may allow for the development of techniques that cannot be accomplished by corresponding CRISPR/Cas9 approaches: By abolishing the nuclease activity of Cas13 while retaining its RNA binding capability, one could use the enzyme to specifically target RNAs to track these transcripts in the cell. Another option would be to fuse Cas13 with different protein domains to affect post-transcriptional processing of target mRNAs, e.g., altering transcript splicing or stability. Specific efforts have been made to investigate these applications with promising results [48–50,52,71].

We were particularly interested in a nuclease-deficient Cas13 variant as a tool to validate specific RNA-protein interactions. For our proof-of-principle approach, we selected the Cas13 variant with the most consistent, robust, and specific interference activity, CasFX4 (hereafter referred to as simply CasFX), and introduced quadruple mutations in the catalytic HEPN domains (R239A/H244A/R858A/H863A). These mutations abolish the nuclease activity but not RNA binding activity in the CasRX variant [50,52,71] (Figure 4A). We first tested whether the mutant CasFX still retained nuclease activity by testing our validated crRNAs against eCFP in the Sg4_CD cell line. As expected, the mutant CasFX failed to interfere with the expression level of eCFP, whereas the wild-type variant worked efficiently (Figures 4B, C). We conclude that this mutant CasFX variant, similar to the corresponding variants in other species, lost its nuclease activity. We hereafter refer this variant as dCasFX (d = dead).

**Figure 4:**
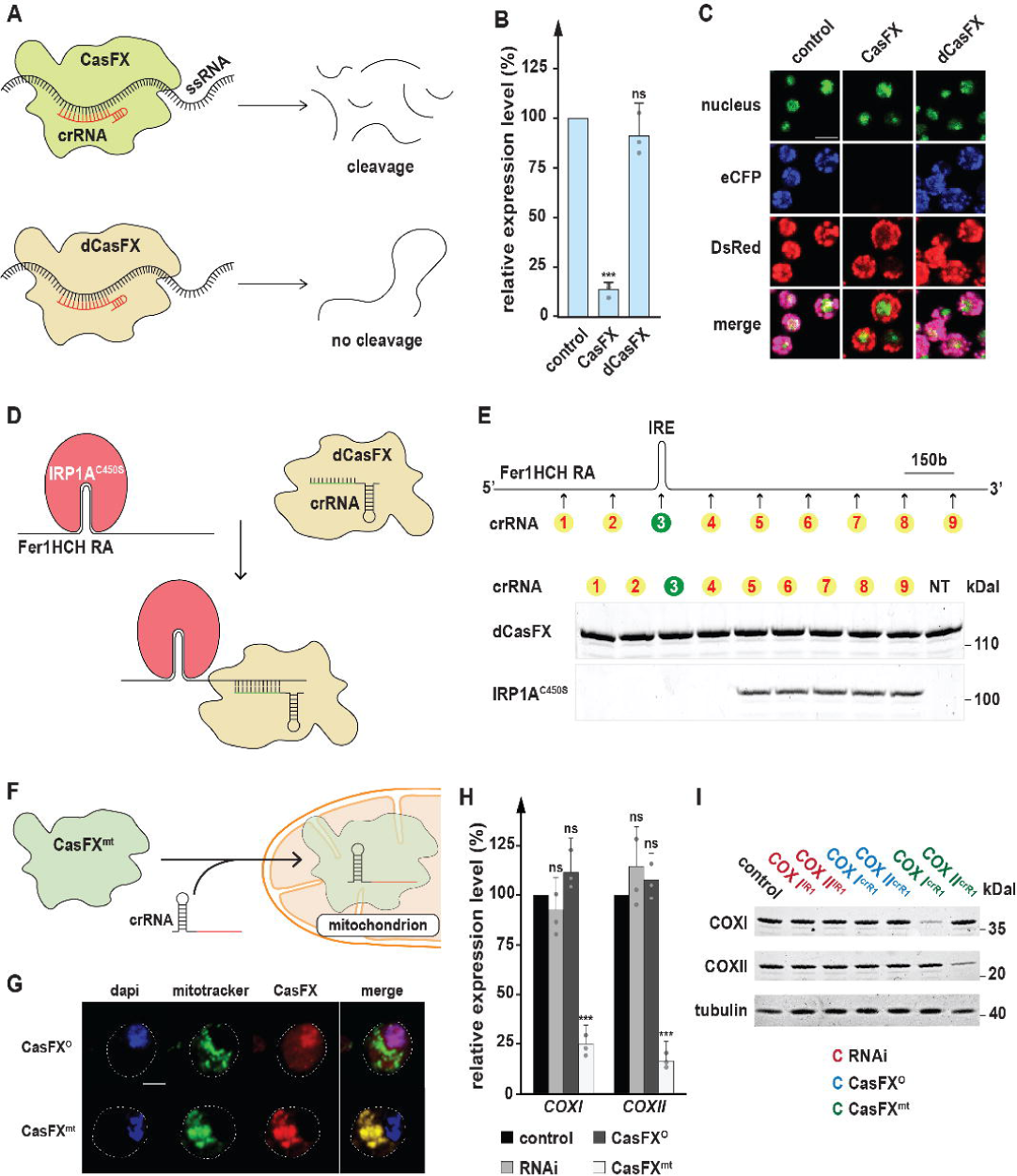
Properties of modified Cas13 variants. **(A)** Schematic of nuclease-dead CasFX (dCasFX) activity. dCasFX carries quadruple point mutations that abolish its nuclease activity. As a result, the dCasFX/crRNA complex can be recruited and bind to target transcripts, but it cannot cleave the RNA. **(B)** Evaluation of Cas13 cleavage efficiency of dCasFX compared to wild-type CasFX. qPCR data represent expression levels of eCFP. Data were normalized to samples treated with blank crRNA (control). * = p-value < 0.05, ** = p-value < 0.01, *** = p-value < 0.001, p-values based on Student’s t-test, error bars represent 95% confidence intervals. **(C)** eCFP fluorescence when targeted by either CasFX or dCasFX. Nuclei were stained with nuclear green DCS1 (Abcam ab138904). eCFP and DsRed fluorescence were measured using their native fluorescence property without using antibody staining. Scale bar = 50 μm. **(D)** Schematic of dCasFX for the validation of RNA-protein interactions. dCasFX and crRNA targeting Fer1HCH-RA mRNA were transfected together in one sample. Fer1HCH-RA and IRP1A^C450S^, a constitutively RNA-binding form of IRP1A that interacts with the iron-responsive element (IRE) in the Fer1HCH-RA mRNA, were transformed together in a different sample. The two samples were each lysed and combined, followed by immunoprecipitation (IP) of dCasFX (utilizing the attached HA tag) to test for the presence of IRP1A in the pull-down assay. **(E)** Western blot showing the IP of dCasFX combined with different crRNAs along *Fer1HCH-RA* mRNA and the detection of IRP1A in corresponding samples. (**F)** Functional schematic of CasFX that carries a mitochondrial localization signal (CasFX^mt^). At the N terminus, CasFX^mt^ is fused with the tim23 mitochondrial signal sequence. Upon binding with crRNA, the complex will localize into mitochondria and target mitochondrial-encoded transcripts. **(G)** Mitochondrial localization of CasFX^mt^. Nuclei were stained with DAPI (blue) while mitochondria were stained with mitotracker green (Cell signaling 9074S) and CasFX polypeptide was stained with anti-HA antibody (red). Scale bar = 25 μm. **(H)** The relative expression level of mitochondrial-encoded transcripts, *COXI* and *COXII*, targeted by RNAi, CasFX^O^, and CasFX^mt^. Data were normalized to samples treated with no transfected plasmid (control). * = p-value < 0.05, ** = p-value < 0.01, *** = p-value < 0.001, ns = not significant, p-values based on Dunnett’s *post-hoc* test, error bars represent 95% confidence intervals. **(I)** Western blotting of COXI and COXII when being targeted by RNAi, CasFX^O^, and CasFX^mt^.

To assess whether crRNA-guided dCasFX would specifically interact in a non-destructive manner with its intended target mRNA, we tested its ability to co-IP a protein known to bind to the same mRNA. As such, immunoprecipitation of dCasFX should pull down the mRNA as well as its bound protein, which can be detected via Western blotting. This approach is useful to validate the RNA-binding activity of dCasFX, as well as the interaction between mRNA and the interrogated protein. To test this, we used an isoform of the ferritin heavy chain 1 mRNA (*Fer1HCH-RA*), which carries a canonical iron-responsive <u>e</u>lement (IRE) at its 5’-end. This IRE allows iron regulatory protein 1A (IRP1A), the *Drosophila* ortholog of human iron regulatory protein 1 (IRP1), to bind to the *Fer1HCH-RA* mRNA [72–76]. Specifically, we used the IRP1A^C450S^ form [74], which is constitutively RNA-binding. We then designed a series of crRNAs that direct dCasFX to its target, *Fer1HCH-RA*, and tested whether immunoprecipitation of dCasFX would also pull down IRP1A. We transfected and lysed cells containing the dCasFX and crRNA components, and mixed this lysate with a second sample obtained by lysing cells containing transfected *Fer1HCH-RA* mRNA and IRP1A^C450S^. By combining the two lysates, the dCasFX/crRNA enzyme should bind to the *Fer1HCH-RA* mRNA/IRP1A^C450S^ complex. If the interaction occurs, immunoprecipitation of dCasFX (via its added HA tag) is expected also to pull down IRP1A^C450S^ (Figure 4D).

A key question for this strategy was how far the recognition site for dCasFX/crRNA needed to be away from the IRE to allow binding of both proteins, dCas13 and IRP1A, to the *Fer1HCH-RA* mRNA. To this end, we generated nine different crRNAs, representing binding sites spaced ∼150 bases apart to roughly cover the entire 1.7 kb *Fer1HCH-RA* mRNA. One of the sites (crRNA #3) partially overlapped with the IRE site, which served as a control to disrupt IRP1A binding. Using this strategy, we found that immunoprecipitation of dCasFX successfully pulled down IRP1A, as long as the cRNA binding site was sufficiently removed from the IRE. As expected, this interaction appeared to be dependent on the distance between the crRNA target site and IRE sequence, since an insufficient distance should cause steric hindrance between the two proteins (Figure 4E). As a control, we used a non-targeting (NT) crRNA to ensure the interactions we observed were specific. The control showed that immunoprecipitation of dCasFX with a non-*Fer1HCH-RA* mRNA-targeting cRNA was not able to pull down IRP1A.

We also tested whether we can simply detect immunoprecipitated *Fer1HCH-RA* mRNA via real-time PCR (qPCR). In the absence of IRP1A, dCasFX appears to bind to the *Fer1HCH-RA* mRNA efficiently, and we found no significant differences between the nine different crRNAs (Figure S5A). Interestingly, when we repeated the experiment in the presence of IRP1A, we noticed a ∼4-fold reduction of immunoprecipitated *Fer1HCH-RA* mRNA when we used cRNAs #1-4 (Figure S5B). This is consistent with the results for co-immunoprecipitated IRP1A (Figure 4E), suggesting that competition between IRP1A and dCasFX (bound to crRNAs #1-4) affected the RNA-binding ability of both proteins. We conclude that dCasFX is a reliable tool to validate interactions between a protein and its candidate target RNA. In addition to RNA-immunoprecipitation, dCasFX could potentially also used for other *in vivo* studies, such as locating a transcript of interest to elucidate its subcellular localization or for co-localization studies, or to determine whether a given protein is bound to its target RNA or unbound.

### Targeting mitochondrial RNAs via Cas13

Like CRISPR/Cas9, Cas13 needs to form a complex with a crRNA before it can identify and cleave its target transcript [22, 23]. Since the Cas13/crRNA complex harbors a single protein, it can be easily tagged with a mitochondrial targeting sequence to cleave RNA in mitochondria, which is not feasible with RNAi. *Drosophila* mitochondria contain multiple copies of circular DNA (mtDNA), which encode tRNAs, rRNAs, and polypeptides important for oxidative phosphorylation. The study of mitochondrial genes is important, because mutations in mtDNA can cause devastating human disorders, such as Leber’s hereditary optic neuropathy, which causes blindness [77,78,79]. To modify CRISPR/Cas13 applications for mitochondrial-encoded transcripts, we added a sequence encoding an N-terminal mitochondrial targeting peptide derived from the nuclear-encoded *translocase of the inner mitochondrial membrane 23* (*tim23*) gene. For this approach, we generated a modified version of our highly efficient CasFX variant, which we termed CasFX^mt^. The CasFX^mt^/crRNA complex is predicted to be imported into the mitochondrial matrix, where it should bind to and cleave the target transcripts (Figures 4F, G).

To test the functionality and efficiency of the CasFX^mt^ variant, we co-transfected CasFX^mt^ with constructs encoding a crRNAs against either *mitochondrial cytochrome c oxidase subunit I* (*mt:CoI*, aka *COXI*) or *mitochondrial cytochrome c oxidase subunit II* (*mt:CoII*, aka *COXII*). Both *COXI* and *COXII* are highly expressed mitochondrial-encoded genes critical for oxidative phosphorylation [80–82]. We analyzed the expression levels of *COXI* and *COXII* via qPCR as well as western blots. To put these results into context, we generated RNAi samples against each of these targets, and used the original CasFX (CasFX^O^, O = original) variant, which lacks the mitochondrial sequence, as a control. In our hands, RNAi targeting either *COXI* or *COXII* had no significant effect on the expression of these two transcripts. Similarly, CasFX^O^/crRNA produced no significant effects (Figures 4H, I). In stark contrast, CasFX^mt^ caused a 4-5-fold reduction of the COX transcripts and resulted in a substantial drop in protein levels as well (Figures 4H, I). To ensure that this result was reproducible, we tested additional RNAi as well as crRNAs sequences, all of which target *COXI* or *COXII* transcripts (Figure S2). In all cases, the observed results were comparable (Figures S5C-E), suggesting that CasFX^mt^ is a useful tool to target mitochondrial-encoded transcripts.

### Cas13-ADAR2 for RNA modification

One intriguing aspect of CRISPR/Cas13 has focused on the modification of RNA, which led to two approaches, namely “<u>R</u>NA <u>e</u>diting for <u>p</u>rogrammable <u>A</u> to I replacement” (REPAIR) and “<u>R</u>NA <u>e</u>diting for <u>s</u>pecific <u>C</u> to <u>U e</u>xchange” (RESCUE) [47, 50]. These methods allow for programmable adenosine-to-inosine editing as well as cytosine-to-uridine editing, respectively. The ability to modify genetic information at the RNA level may be advantageous, because, unlike Cas9 which causes a permanent change in the genome, RNA modifications via Cas13 are reversible due to RNA turnover [8,12,39,74]. As such, Cas13-based approaches may be suitable for future therapies, where Cas13 could be used to repair missense mutations in transcripts without affecting a patient’s genome.

In the REPAIR systems used in mammalian cells, the nuclease-dead PspCas13b was fused to the RNA-modifying domain of Adenosine Deaminase Acting on RNA 2 (ADAR2). In their original approach, Cox et al. found that the first REPAIR version (REPAIRv1) had substantial off-target activity. Subsequently, they generated REPAIRv2, which harbored two point mutations in the ADAR2 domain (T375G and E488Q). This version showed high specificity and robustness in mammalian cells [50].

Given its success in mammalian cell systems, we wondered whether a Cas13-ADAR fusion would be functional in *Drosophila*. The insect ADAR protein appears to function similarly to its human counterpart [83], suggesting that constructs based on mammalian ADAR2 would work in *Drosophila*. We first fused the above-described dCasFX to the mutant human ADAR2 domain that carries equivalent mutations as the REPAIRv2 we mentioned earlier. We refer to this construct as FREPAIRv2 (F = fruit fly), and tested for its editing efficiency (Figure 5A). To test for Cas13-ADAR2 activity, we generated a system that uses a dual reporter transgene in the *Drosophila* embryo cell line Sg4-PP-27F. Similar to the earlier described Sg4_CD line; this cell line carries the independently expressed *eCFP* and *DsRed* transcription units in the genome, each with their own *actin5* promoters. However, unlike the Sg4_CD line, we introduced a point mutation into the eCFP coding region that converts a tryptophan residue 57 (W57*) TGG into an early stop codon (TGA), which we refer to as eCFP*. Also, we termed this new cell line “Sg4*” line to distinguish it from the original Sg4_CD (Figure S1B). Next, we co-expressed FREPAIRv2 and an eCFP-crRNA, which carries a single mismatch A to C at the position that corresponds to the introduced stop codon (Figures 5A, B). If the FREPAIRv2 is capable of editing its target RNA encoded by *eCFP\**, the stop codon should be reverted to the wild-type tryptophan residue (W57), and the resulting full-length eCFP should be detectable via Western blotting and, if efficiency is sufficiently high, via fluorescence from the restored CFP. Using this strategy, we found that we were able to detect fluorescence at a wavelength of 405 nm as early as 36 hours after transfection, indicating that detectable levels of eCFP had been produced. eCFP fluorescence continued to increase, with substantially higher levels at the 60-hour time point (Figure 5D). When we conducted Western Blots to validate these data, we saw corresponding results, with detectable eCFP protein at 36 hours and progressively higher levels from 42 to 60 hrs after transfection (Figure 5C). We conclude that Cas13-ADAR2 works effectively in *Drosophila* and can be used to modify target mRNAs, such as reverting transcripts carrying missense mutations without altering the genome.

**Figure 5:**
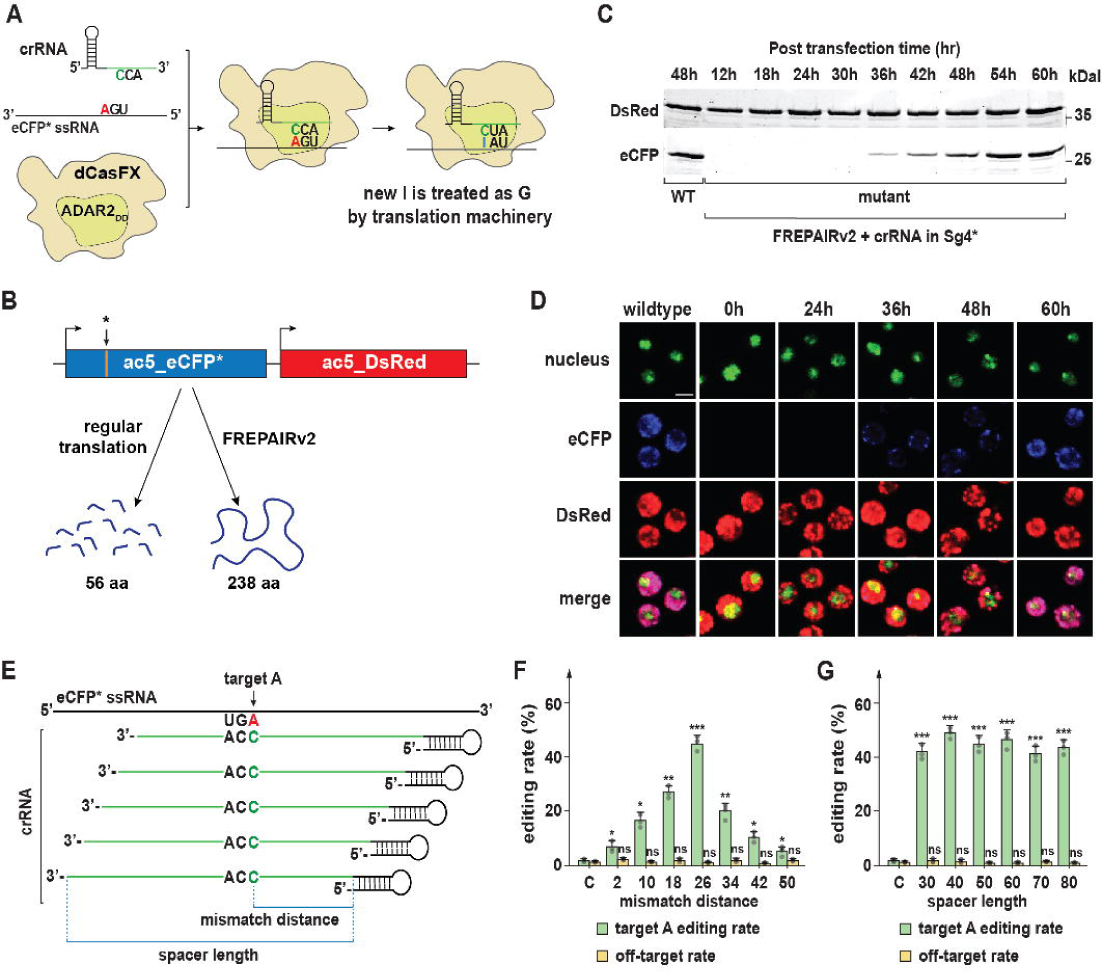
Adaptation of the REPAIRv2 system to modify RNA in *Drosophila* Sg4 cell culture. **(A)** Schematic for the *Drosophila*-modified REPAIRv2 system (FREPAIRv2), to modify a mutant eCFP transcript. Mutant eCFP carries an early stop codon that normally encodes Tryptophan at residue 57 (W57*). By generating an A to C mismatch in the crRNA spacer that corresponds to the stop codon, the ADAR2_DD_ domain will change the equivalent adenosine (A) to inosine (I). Inosine will be treated as guanosine by the translation machinery. **(B)** Schematic of FREPAIRv2 outcome. Originally, the mutant eCFP transcript harbors a stop codon at position 57, which will generate a short polypeptide with 56 amino acids. However, once modified by FREPAIRv2, codon 57 will be reverted to wild-type tryptophan, and restore the production of a full-length polypeptide. **(C)** Western blotting monitoring eCFP productions relative to transfection time. **(D)** Fluorescence emitted by eCFP relative to transfection time. Nuclei were stained with nuclear green DCS1 (Abcam ab138905). eCFP and DsRed fluorescence were measured based on their natively emitted fluorescence. Scale bar = 50 μm. **(E)** Schematic of crRNAs that we used for FREPAIRv2. We considered two criteria for the crRNA design: i) mismatch distance from the first nucleotide and ii) spacer length. **(F)** Editing rate and off-target rate of FREPAIRv2 concerning mismatch distance when spacer length was kept at a constant 50 nucleotides. Error bars represent standard deviation. **(G)** Editing rate and off-target rates of FREPAIRv2 in relation to spacer lengths when the mismatch distance was kept at the constant position 26. Error bars represent standard deviation.

For the above approach, we followed a similar path that was used in the original study [50] where the mismatch (C→A) was placed in the center of the crRNA spacer, measured at the 26^th^ nucleotide of 50 nucleotides (nt) spacer, relative to the stem loop-forming direct repeat of the crRNA. To evaluate the editing efficiency in correlation to mismatch position and spacer length, we tested a series of crRNA constructs with the same spacer length of 50 nt; however, we changed the relative mismatch distance to the hairpin by increments of 8 nt (Figure 5E). We then performed reverse transcription and sequenced a minimum of ten randomly selected eCFP cDNAs per construct. This was followed by sequencing to assess the fraction of clones that harbored the repaired codon for tryptophan #57, expressed as editing rate (Figure 5F). Based on our findings, the crRNA that carried the mismatch at position 26 relative to the hairpin (“mismatch distance”, Figure 5E) resulted in the highest efficiency (Figure 5F), consistent with other studies [50]. We then tested the effect of varying spacer length while keeping the mismatch distance at 26 nt. We tested spacer lengths from 30 nt to 80 nt, and in all cases, we observed similar efficiencies, all of which were comparable to a 50 nt spacer (Figure 5G). Based on these findings, we conclude that FREPAIRv2 works best when using a mismatch distance of 26 nt, whereas the spacer length did not appear to affect the editing efficiency [50].

To evaluate the off-target tendencies of FREPAIRv2 in *Drosophila* cells, we examined the cDNA sequences for additional A→I modifications, which is straightforward since inosine is recognized as guanosine by the reverse transcriptase. However, we scored any unpredicted sequence deviations as potential off-target events and plotted them relative to the mismatch distances and spacer lengths (Figures 5F, G). This strategy revealed that some off-target effects persisted, albeit at a low level across all crRNAs that we tested. Given that these effects are random, and distributed across multiple RNA molecules, it appears likely that this off-target activity has no or inconsequential impact on phenotypes. However, future studies are needed to improve the specificity of this editing system further.

### Generation and characterization of transgenic Cas13 flies

Our data demonstrated that Cas13 works well in *Drosophila* Sg4 cells and can be used for purposes beyond RNA cleavage. We next sought to generate transgenic fly lines carrying Cas13 variants and characterize their efficacy *in vivo*. To this date, no study has analyzed the usability Cas13 in live organisms to the best of our knowledge. As such, it is critical to establish whether Cas13-based technology is suitable for *in vivo* studies. Furthermore, we were interested in creating a system that allows for temporal and spatial control over Cas13 expression. To this end, we have previously created a *Drosophila* toolkit for CRISPR/Cas9 based on Gateway-compatible cassettes that allow researchers to insert specific enhancers that drive the expression of the Cas transgene in a tissue of interest [8, 59]. While this generates more upfront work compared to Gal4/UAS-based systems driving the expression of Cas9 [12, 37], it does simplify the downstream workflow. Also, it reduces unspecific effects since one requires fewer transgenes to build the necessary fly genotype. We, therefore, decided to create a similar Cas13 toolkit. In total, we manufactured two general Cas13 vectors, one based on CasFB and one that uses CasFX, both of which displayed the highest catalytic efficiency in Sg4_CD cells. For our *in vivo* strategy, we limited our efforts to constructs that would interfere with RNA expression (Figure S6A). Based on these all-purpose vectors, we then generated four transgenic lines for further characterization, named here *act_CasFB*, *UAS-CasFB*, *act_CasFX*, *UAS-CasFX* (Figure S6A).

For the generation of crRNAs, we used the previously described multiplexed pCFD5 vector and implemented changes suitable for Cas13 crRNA processing [12]. We refer to the new plasmids as i) pC13B, which expresses CasFB-compatible crRNAs under control of the U6:3 promoter and ii) pC13X, which expresses CasFX-compatible crRNAs under control of the U6:3 promoter (Figures S6B, C). Both plasmids will ubiquitously express the tRNA:crRNA units. As the tRNA is processed, its cleavage will result in the release of mature crRNAs that form complexes with Cas13 enzymes. The cloning procedures for these new crRNA plasmids are overall similar to those for the pCFD5 vector, but, since some differences exist, we include a detailed protocol in the supplementary material (see supplemental method S1).

To evaluate the efficiency of our transgenic Cas13 constructs *in vivo*, we generated seven transgenic crRNAs targeting three genes that we study in our lab. This includes *phantom* (*phm*) and *disembodied* (*dib*), two well-characterized genes involved in ecdysone synthesis in *Drosophila* [84, 85] as well as the third gene, *Iron Regulatory Protein 1A* (*IRP1A*), a gene critical for cellular iron homeostasis [74, 86]. Classic mutants of *phm* and *dib* display embryonic lethality while *IRP1A* mutant animals die as first instar larvae (L1) [8,59,74,84,87]. In contrast, using PG-specific somatic CRISPR/Cas9 strategies, *phm^gR^* (gRNA for CRISPR Cas9) caused L1 arrest, while *dib^gR^* and *IRP1A^gR^* both caused third instar (L3) larval arrest (Figures 6A-C) [8,59,74]. In addition, PG-specific disruption of *IRP1A* via somatic CRISPR/Cas9 caused a porphyria-like phenotype due to iron deficiency (Figure 6D) [74].

**Figure 6:**
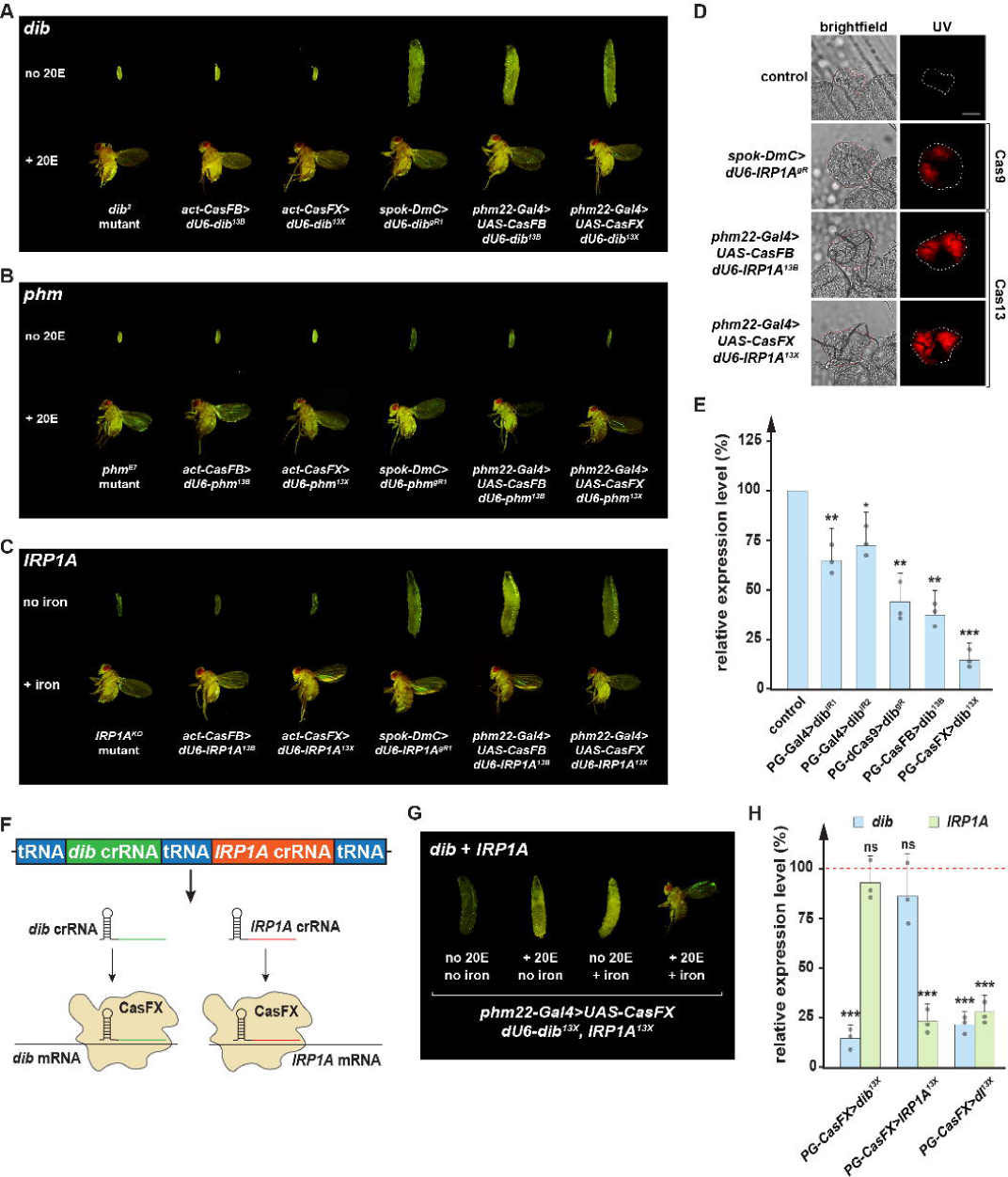
Efficiency of *Drosophila* codon-optimized CRISPR/Cas13 *in vivo*. **(A)** Comparison of phenotypes from a classic *disembodied* mutant (*dib^2^*), ubiquitous knock-down of *dib* via CasFB/*dib*^13B^, CasFX/*dib*^13X^, prothoracic gland-specific manipulation via CRISPR/Cas9, or Cas13 of *dib* in the presence or absence of 20-Hydroxyecdysone (20E). **(B)** Comparison of phenotypes from a classic *phantom* mutant (*phm^E7^*), ubiquitous knock-down of *phm* via CasFB/*phm*^13B^, CasFX/*phm*^13X^, PG-specific manipulation via CRISPR/Cas9, or Cas13 of *phm* in the presence or absence of 20-Hydroxyecdysone (20E). **(C)** Comparison of phenotypes from a classic *iron regulatory protein 1* mutant (*IRP1A^KO^*), ubiquitous knock-down of *IRP1A* via CasFB/*IRP1A*^13B^, CasFX/*IRP1A*^13X^, PG-specific manipulation via CRISPR/Cas9, or Cas13 of *IRP1A* in the presence or absence of iron in the diet. **(D)** Porphyria phenotype in PG-specific *IRP1A* knock-down. Scale bar = 250 μm. **(E)** Relative *dib* expression levels in samples representing different PG-specific loss-of-function strategies, including RNAi (IR), dCas9-mediated transcriptional interference, and Cas13 cleavage. Ring glands were dissected from larvae at 42-hour after the L2/L3 molt. * = p-value < 0.05, ** = p-value < 0.01, *** = p-value < 0.001, ns = not significant, p-values based on Dunnett’s *post-hoc* test, error bars represent 95% confidence intervals. **(F)** Schematic of a construct containing two crRNAs simultaneously targeting *dib* and *IRP1A* mRNA. **(G)** Comparison of phenotypes from PG-CasFX/*dI*^13X^ in the presence or absence of either 20E, iron, or both. **(H)** Relative expression levels of *dib* and *IRP1A* in single or double knock-down PG samples. Data were normalized to the expression of these genes in controls. Ring glands were dissected from larvae at 42-hour after the L2/L3 molt. * = p-value < 0.05, ** = p-value < 0.01, *** = p-value < 0.001, ns = not significant, p-values based on Student’s t-tests, error bars represent 95% confidence intervals.

When we crossed the Cas13-compatible crRNAs (referred to as 13B for CasFB-compatible cRNAs and 13X for CasFX-compatible crRNAs) targeting either *phm*, *dib* or *IRP1A* with either ubiquitously expressed or PG-specific Cas13 variants, we observed the same developmental defects we found with our previous strategies (Figures 6A-D, S2, S7, Table S3), indicating that Cas13 worked effectively in *Drosophila*. The fact that *phm^13B^, phm^13X^, dib^13B^, and dib^13X^* individuals were rescued to adulthood when reared on 20E-supplemented media [8, 59], and that *IRP1A^13B^*, as well as *IRP1A^13X^* animals, reached adulthood when dietary iron was provided [74], strongly suggested that the activity Cas13 was highly specific (Figures 6A-C, S7).

In addition to the above phenotypic analysis, we evaluated *dib* expression levels via qPCR. We compared the results to other tissue-specific loss-of-function techniques, including samples from two independent RNAi lines and samples from one line where we used transcriptional interference via dead Cas9 (dCas9) to target *dib*. We found that the two RNAi lines reduced dib expression by 30-40%, whereas the CRISPRi approach via dCas9 lowered *dib* expression by 50-60%. Concerning the new Cas13 lines, CasFB reduced *dib* expression by 55-65%, equivalent to the dCas9 data. Remarkably, CasFX showed the strongest knock-down, and robustly reduced *dib* expression by 80-90% (Figure 6E). These data indicated that Cas13 transgenes work *in vivo* and may exceed the efficacy of other techniques.

We also tested the ability to target multiple transcripts with a single transgene. For this, we used the pC13X vector and generated a dual-crRNA transgenic line (termed dI^13X^) that ubiquitously expressed a crRNA targeting *dib* mRNA as well as a crRNA targeting the *IRP1A* transcript (Figure 6F). Target sites for either of these transcripts were the same as before (Figures 6A, 6C, S2). As expected, the animals arrested development at the L3 stage, similar to targeting the *dib* and *IRP1A* transcripts individually. Consistent with this, neither 20E- nor iron-supplementation alone could rescue these double knock-downs, however, a diet supplemented with both 20E and iron caused a significant rescue to adulthood (Figure 6G). This makes sense since the two cRNAs interfered with ecdysone production and the regulation of cellular iron homeostasis. To assess whether the simultaneous knock-down of two genes was as efficient as targeting these genes individually, we evaluated *dib* and *IRP1A* expression levels via qPCR. We found no significant difference in any of these approaches suggesting that there is no penalty when targeting two genes at the same time (Figure 6H).

## Discussion

### RNA degradation efficiency of Cas13 in *Drosophila*

We evaluated eleven variants of each reported Cas13 ortholog in *Drosophila* Sg4 cells, including the well-characterized variant from the original studies and ten *Drosophila*-optimized variants. Among all Cas13 enzymes tested, we observed a wide range of efficiencies, even between the versions from the same ortholog. Among them, CasRX and its *Drosophila*-optimized variants CasFX appeared to have the highest efficiency. For the Cas13a and Cas13b variants, we also identified the optimized variants with reliable efficiency. Even though they were less efficient than CasFX, these variants may still prove useful in circumstances where only a moderate knock-down is desired. On the other hand, Cas13c variants did not significantly alter the expression of target transcripts. We hypothesize that this was caused by several factors: (i) Cas13c is the least characterized Cas13 enzyme, and it might use a mechanism that differs from the other Cas13 enzymes. (ii) Even though the low efficiency of Cas13c was in agreement with previous studies conducted in other species, we cannot rule out the possibility that the Cas13c variants we used were not ideally suited for *Drosophila*, and (iii) Cas13c might still require a PFS for optimal activity in the fruit fly. Future studies will need to address this.

We noticed that the expression of the PspCas13b and CasRX variants resulted in considerable toxicity when animals were homozygous for these transgenes, causing lethality during the first (L1) or second (L2) instar larvae (Figure S8). Interestingly, animals heterozygous for PspCas13b and CasRX transgenes showed no significant lethality. In contrast, animals homozygous for our *Drosophila*-optimized Cas13 transgenes, namely CasFB and CasFX, showed only moderate lethality, with 51% to 58% reaching adulthood, respectively (80-85% is expected in wild type populations). As expected, animals heterozygous for these transgenes appeared normal (Figure S8). The lethality of Cas13 transgenic animals was also reported in a recent study [88], similar to the results of early versions of Cas9 in *Drosophila* [8, 10]. Since we observed a wide range of efficiencies between the variants, it is possible that each variant also exhibits different levels of toxicity. While the reasons for the relatively high lethality of the original PspCas13b and CasRX constructs (in a homozygous setting) remain unclear, our data suggest that each variant is unique and that perhaps using codon-optimized versions help to reduce the toxicity associated with Cas13.

### Beyond RNA cleavage

A few studies have shown that Cas13 may be useful in a broad range of applications, and not just RNA cleavage. In this study, we have demonstrated that dCasFX can validate RNA-protein interactions by using an appropriately designed crRNA. We also showed that by adding a mitochondrial localization sequence, one could recruit the CasFX^mt^/crRNA complex into mitochondria and target mitochondrial-encoded transcripts. We also adopted the REPAIRv2 system from mammalian cell culture into *Drosophila* Sg4 cells and showed that this system, FREPAIRv2, can efficiently modify target transcripts with an overall low off-target rate. We have not tested other potential applications; however, in theory, Cas13 can be modified for many approaches to study RNA, including splicing, transcript stabilization, or RNA localization.

Cas13 may have far-reaching implications for simplifying diagnostics. Recently, the outbreak COVID-19 caused by SARS-CoV-2 has resulted in a global health threat. To develop a fast test for COVID-19, the <u>s</u>pecific <u>h</u>igh-sensitivity <u>e</u>nzymatic reporter un<u>lock</u>ing (SHERLOCK) protocol, a recently developed Cas13-based diagnostic test for infectious diseases, can detect the virus in 50 min [89, 90] (https://mcgovern.mit.edu/2020/02/14/enabling-coronavirus-detection-using-crispr-cas13-an-open-access-sherlock-research-protocol/). In an independent study, CRISPR/Cas13 was also used to detect SARS-CoV-2 [91]. Together, these studies demonstrate the enormous potential of Cas13 as a diagnostic and therapeutic tool.

### From in vitro to in vivo

A significant part of the work presented here was based on cell culture experiments. These approaches were ideal to economically evaluate the efficiencies of multiple Cas13 versions in *Drosophila*. However, our ultimate goal is to establish CRISPR/Cas13 approaches for *in vivo* studies in model organisms, which has not been accomplished yet. Based on our results of transgenic CRISPR/Cas13 flies, CasFX and CasFB can efficiently target and cleave transcripts of interest *in vivo*, and as such, represent a compelling alternative to existing methods. This study may also help scientists working with other model organisms to optimize their approach for implementing Cas13 *in vivo*.

### The CRISPR/Cas13-based toolkit

This study has generated two collections of Cas13/crRNA toolkits to study in either cell culture or organisms. For the cell culture toolkit, we have produced the pC13cr01 vectors, which allow the co-transfection of Cas13 variants and the crRNA corresponding to the target transcript. With this vector, one only needs to digest the crRNA backbone with the BbsI enzyme and clone the target site for the crRNA, similar to the generation of the Cas9-compatible gRNA system in pCFD5 or pCFD6 plasmids. For *in vivo* work, we also established a similar system with Cas13 transgenes already available from our study. Researchers will need to generate their crRNAs against the target transcript. For this, we provide the pC13B and pC13D vectors with the same cloning procedure as pCFD5. We also provided a supplemental method section with a detailed description of the cloning procedures. On the other hand, the UAS-based versions of Cas13 transgenes will also allow scientists to spatially and temporally manipulate Cas13 activity and study transcript of interest at desired tissues.

### Conclusions

Just like CRISPR/Cas9 allows for the manipulation of DNA, Cas13 enables us to target any transcript of interest. This is beneficial for approaches where researchers do not want to alter the DNA of the gene of interest, since Cas13 controls gene expression on the RNA level, similar to RNAi. Furthermore, current evidence suggests that Cas13, especially variants from the Cas13d family, display minimal off-target tendencies, and this might help quell concerns regarding RNA targeting. Even though it might be too early to make conclusions about the off-target activity of Cas13, we believe that its high specificity holds excellent promise for future applications. Also, the ability to modify Cas13, such as targeting Cas13 to mitochondria, further expands the range of future applications for this methodology.

## Methods

### Generation of *Drosophila* optimized Cas13 orthologs (DmCas13)

To generate fruit fly codon-optimized Cas13 variants, the original Cas13 nucleotide sequences were evaluated by using two independent web tools: i) ATGme (https://atgme.org) and ii) OPTIMIZER (http://genomes.urv.es/OPTIMIZER) [92, 93] with the customized codon usage frequency specific for *Drosophila* [94–96]. The two indices, namely the Codon Adaption Index (CAI) and the Effective Number of Codons (ENC) were used to obtain the optimized sequences. CAI has value ranges from 0 to 1 and is used to evaluate the similarity between codon usage of a gene and codon usage of the reference group [97]. Thus, at least in theory, the higher the CAI value, the higher is gene expression [98, 99]. On the other hand, ENC is a measure of codon usage bias with values between 20 and 61. Since the expression of a gene is usually dependent on the availability of tRNA species, one would expect that genes with higher expression will use a smaller subset of codons recognized by the most abundant tRNAs, resulting in lower ENC values [100]. Taking these two factors into consideration, we picked the top 10 variants per Cas13 subtype for further investigation (Table S1). We reasoned that it would not suffice just to choose the top-scoring variant, and therefore, we also selected other high-scoring sequences. We generated the selected variants via a combination of mutagenesis of the original Cas13 sequences and fusing gBlocks gene fragments from Integrated DNA Technologies (IDT) (Table S4).

### Design and generation of target crRNAs

The very first Cas13 proteins that were characterized in bacteria required a sequence constraint, the PFS, to ensure target cleavage efficiency. This includes *Leptotrichia shahii* Cas13a (LshCas13a), *Bergeyella zoohelcum* Cas13b (BzoCas13b) and *Prevotella buccae* Cas13b (PspCas13b) [50, 53]. However, further investigation of PspCas13b in mammalian and plant and other Cas13 orthologs showed high target RNA degradation efficiencies even in the absence of PFS [23,46,49,52]. While this gives researchers some flexibility over target site selection, it is necessary to consider the secondary structure of target transcripts, since this negatively affected knock-down efficiency [23, 50]. To assess secondary structures, we used two independent online tools, namely RNAfold (http://rna.tbi.univie.ac.at/cgi-bin/RNAWebSuite/RNAfold.cgi) and RNA structure (https://rna.urmc.rochester.edu/RNAstructureWeb/Servers/Predict1/Predict1.html) [101–104]. Besides, we also used the siRNA design tool RNAxs (http://rna.tbi.univie.ac.at/cgi-bin/RNAxs/RNAxs.cgi) to find the regions of transcripts with good accessibility to narrow down the target region space for designing gRNAs [105]. For the case of Cas13a orthologs, we compared the target sequences with the online CRISPR-RT tool (http://bioinfolab.miamioh.edu/CRISPR-RT/interface/C2c2.php) [106]. The crRNA cassette was amplified and cloned into a pre-digested expression vector backbone via the Gibson reaction. All crRNAs used in this study were driven by the *Drosophila* U6:3 promoter (dU6:3). For more information regarding crRNA cloning, see supplement method S1.

### Generation of transfection plasmids

For a list of plasmids, we generated for this study, see supplemental table S2. The original plasmids we used for this project were obtained from Addgene: pCFD3 (#49410), pCFD5 (#73914), pACG:eCFP (#32597), pDsRed-attP (#51019), Ac5-Stable2-Neo (#32426), pC0056-LwaCas13a-msfGFP-NES (#105815), pC0040-LwaCas13a crRNA backbone (#103851), pC0046-EF1a-PspCas13b-NES-HIV (#103862), pC0043-PspCas13b crRNA backbone (#103854), pC0054-CMV-dPspCas13b-longlinker-ADAR2DD (E488Q/T375G) (103870), pXR001: EF1-CasRX-2A-eGFP (#109049), pXR004: CasRX pre-gRNA cloning backbone (#109054), pBID-UASc (#35200), [10,12,23,35,46,50,107–110]. We also obtained plasmids from the *Drosophila* Genetic Resource Center (DGRC): pAFW (#1111), pAHW (#1095), act-PhiC31-integrase (#1368). We also used plasmids we previously generated, enDmC, to generate some constructs for this study [8]. pMT-Gal4-puro plasmid was a kind gift from Christoph Metzendorf (University of Uppsala). All fragments used for the cloning step were amplified via PCR using Q5 high fidelity DNA polymerase (NEB #M0491S) (table S4) and fused together via Gibson assembly reaction [111].

### Generation of transgenic cell lines

The original Sg4-PP-27F (#238) cell culture line was obtained from *Drosophila* Genetics Resource Center (DGRC) and grown in the HyClone SFM4 Insect cell culture (SFM4) medium (GE Lifesciences SH30913.02) with 1% (v/v) Streptomycin-Penicillin (Sigma P4333) following standard procedures (Invitrogen). To generate the transgenic Sg4_CD cell line, Sg4-PP-27F cells were co-transfected with two different plasmids, where one plasmid contained the PhiC31 integrase gene, and the other was the dual-reporter plasmid (Figure S1A). The dual-reporter transgenic construct also contained a *Neo^R^* gene, which allows for resistance to Geneticin G418 (Sigma 4727878001). 48-72 hours after co-transfection, cells were washed and grown in SFM4 medium supplemented with G418 at the final concentration of 200 μg/ml. Transfected cells were maintained on this type of medium (SFM4 with 1% Streptomycin-Penicillin and 200 μg/ml G418) for at least four passage rounds before being tested for the integration of transgenic constructs via Sanger sequencing.

### DNA extraction from cells

Cells were grown, and DNA was extracted as previously described [59]. In brief, cells were collected as pellets and filled with 20 μl of DNA extraction buffer (10 mM Tris-HCl pH 8.2, 25 mM NaCl, 1 mM EDTA pH 8.0, 0.2% v/v Triton X-100, 1x proteinase K (AM2546)). The mixture was vortex for 3x 30s and incubated at 37°C for 30 minutes before heat-inactivated at 95°C for 5 minutes. Cell lysates were centrifuged at 12,000 x g at 4°C for 5 minutes, and the supernatant was transferred to a new collection tube. 1 μl of supernatant was used for PCR amplification at the genomic region spanning target sites. PCR products were purified using the HighPrep^TM^ PCR reagent from MagBio (AC-60005) following the manufacturer’s protocol.

### Cell culture transfection

Cells were grown in SFM4 medium with 1% streptomycin-penicillin, 200 μg/ml G418, and transfected by the Calcium Phosphate-based method (Invitrogen). To study the effects of different Cas13 variants, puromycin was added to medium on the second day after transfection at the final concentration of 5 μg/ml, similar to what was previously used [112, 113]. Cells were collected seven days after transfection to ensure the turnover of already translated eCFP polypeptides [114]. Transfected cells were washed in ice-cold 1x PBS for 3x 5 min and collected for later experiments.

### Cell immunostaining

On the first day of the transfection experiment, coverslips were pre-cleaned in 70% ethanol and placed into a transfection plate (Sigma CLS3516). Cells were then seeded and transfected following the standard procedures (Invitrogen). This allows cells to adhere to the coverslips for subsequent immunostaining. Subsequent transfection procedures were carried out as described in the cell culture transfection section. Seven days after transfection, coverslips were transferred to a clean transfection plate for immunostaining, while cells in the supernatants were collected for cell lysis and protein extraction.

Samples were fixed in 1x PBS 4% formaldehyde (ThermoFisher #28906) for 15 min at room temperature (RT) with gentle shaking followed by washing in 1x PBS 0.3% Triton (Sigma #T9284) (PBS3T) for 3x 10 min. Samples were blocked at RT for 30 min in blocking solution (1x PBS3T 5% normal goat serum (Abcam ab138478)) and incubated in primary antibody dilution buffer (antibody diluted in 1x PBS3T and 1% BSA) for 1 hour at RT. Samples were then washed in 1x PBS3T for three times with 10 min each, incubated in secondary antibody dilution buffer for 1 hour at RT, and then washed in 1x PBS3T with either 1:50,000 DAPI (Cell Signaling #4083) or 1:2,000 Nuclear Green DCS1 (Abcam ab138905) for 3x 10 min. Samples were mounted in Vectashield mounting medium (#VECTH1000). Pictures were taken on Nikon Eclipse 80i Confocal C2+ microscope/camera. We used the following reagents: a monoclonal mouse anti-HA-tag antibody (Abcam ab18181) at the ratio of 1:1000 for 3xHA tagged Cas13 orthologs, mitotracker green (Cell signaling 9074S) at the concentration of 400n M for staining mitochondria, monoclonal mouse anti-MTCO1 (Abcam ab14705) at the ratio of 1:2000 and monoclonal rabbit anti-MTCO2 (Abcam ab79793). Secondary antibodies were obtained from Abcam and used at the ratio of 1:2000 ratio, including goat anti-mouse IgG H&L Alexa Fluor 488 (ab150113), goat anti-mouse IgG H&L Alexa Fluor 555 (ab150114), goat anti-rabbit IgG H&L Alexa Fluor 555 (ab150078). eCFP and DsRed signals were captured based on their fluorescence properties without antibody staining. For quantification of the eCFP signal, the mean pixel values of the images were analyzed using ImageJ as the corrected total cell fluorescence (CTCF) following the formula: CTCF = selected cell intensity – (area of the chosen cell * background intensity). The CTCF values were averaged from all biological replicates and normalized to the normalized average CTCF values of no-targeting (NT) crRNA samples.

### Western blotting of cell extracts

For cell lysis and western blotting, 7-day post-transfection cells were collected by centrifugation at 1,000 x g for 10 min at 4°C and supernatant was removed as much as possible. Cells were washed in ice-cold 1x PBS for 3 × 10 minutes and lysed in 90 μl lysis buffer (1x PBS, 1% Triton, 1x proteinase K inhibitor) by vortexing for 15 seconds every 10 min for up to 1 hour. Cell lysate was mixed with fresh 4x Laemmli buffer (0.25 M Tris pH 6.8, 8% SDS, 40% Glycerol, 25% β-mercaptoethanol, 0.2% bromophenol blue) at the ratio of 3:1 (v/v). 40 μl of the mixture (1/3 total volume) was loaded on 12.5% SDS gel. Later steps, including gel electrophoresis and western blotting, were carried out following the manufacturer’s (Abcam) instructions. To detect eCFP, monoclonal rabbit anti-GFP-tag antibodies (Invitrogen G10362) were used at a ratio of 1:1000, followed by incubation with a goat anti-rabbit IgG H&L HRP secondary antibody (Abcam ab97051) at a ratio of 1:20,000. To detect DsRed, monoclonal mouse anti-DsRed antibody (Santa Cruz sc-390909) was detected at the ratio of 1:1000, followed by incubation with a goat anti-mouse IgG H&L HRP secondary antibody (Abcam ab97023) at the ratio of 1:20,000. To detect COXI and COXII, monoclonal mouse anti-MTCO1 antibody (Abcam ab14705) and monoclonal rabbit anti-MTCO2 antibody (Abcam ab79393), respectively, were used at a ratio of 1:500. To detect tubulin, which served as a loading control, monoclonal mouse anti-β-tubulin antibodies (Sigma 05-661) were used at the ratio of 1:10,000. Blots were scanned for image acquisition with a ChemiDoc imaging system (Bio-Rad), and bands intensity was measured using ImageJ.

### Nuclease-dead dCasFX-IRP1A^C450S^ co-immunoprecipitation

The dCasFX/crRNA complex and IRP1A^C450S^/Fer1HCH RA cDNA was transfected independently. In one sample, dCasFX and the crRNA corresponding to the Fer1HCH-RA transcript were cloned into the same plasmid pC13cr01 (Figure S1D), while in another approach, IRP1A^C450S^ and Fer1HCH RA cDNA were similarly cloned into the same plasmid as pC13cr01. IRP1A^C450S^/Fer1HCH-RA co-transfection was carried out at a 10x higher ratio compared to each dCasFX/crRNA transfection alone. Transfected samples were lysed using 200 μl lysis buffer (1x PBS, 1% Triton, 1x proteinase K inhibitor) by vortexing for 15 seconds every 10 min for up to 1 hour. Lysates of samples transfected with IRP1A^C450S^/Fer1HCH-RA were combined and evenly distributed among ten groups of dCasFX/crRNA lysate. This ensured that each lysate had a similar amount of IRP1A^C450S^/Fer1HCH-RA complex. The mixed lysate was incubated with pre-crosslinked HA Dynabeads protein G (Invitrogen 10004D) following the manufacturer’s directions. Samples were eluted in 4x Laemmli buffer (0.25 M Tris pH 6.8, 8% SDS, 40% Glycerol, 25% β-mercaptoethanol, 0.2% bromophenol blue).

### *Drosophila* stocks and husbandry

We obtained the following stocks from the Bloomington *Drosophila* Stock Center: *w^1118^* (#3605), *dib^2^/TM3 Sb^1^* (#2776), *phm^E7^/FM7c* (#2208), *y^1^v^1^P[nos-PhiC31.NLS]X; attP40(II)* (#25709), *y^1^v^1^P[nos-PhiC31/int.NLS]X; attP2(III)* (#25710). Stocks *UAS-dib*-RNAi (1) (#101117), *UAS-dib*-RNAi (2) (#16827), *UAS-phm*-RNAi (#108359) were obtained from the Vienna *Drosophila* Resource Center. *y^2^cho^2^v^1^* (TBX-0004), *y^2^cho^2^v^1^; sco/CyO* (TBX-0007), *y^2^cho^2^v^1^/Y^hs-hid^; Sp/CyO* (TBX-0008), *y^2^cho^2^v^1^; Sp hs-hid/CyO* (TBX-0009), *y^2^cho^2^v^1^; Pr Dr/TM6C, Sb Tb* (TBX-0010) were obtained from the National Institute of Genetics of Japan (NIG). *act_DmCas13B/CyO GFP*, *UAS-DmCas13B*, *act_DmCasRX/CyO GFP*, *UAS-DmCasRX*, *y^1^v^1^;P[pCFD5 dib.KO dgRNA]attP40* (dU6-dib^gR13B^), *y^1^v^1^;P[pCFD5 dib.KO dgRNA]attP40* (dU6-dib^gR13D^), *y^1^v^1^;P[pCFD5 dib.KO dgRNA]attP40* (dU6-phm^gR13B^), *y^1^v^1^;P[pCFD5 dib.KO dgRNA]attP40* (dU6-phm^gR13D^), *y^1^v^1^;P[pCFD5 dib.KO dgRNA]attP40* (dU6-IRP1A^gR13B^), *y^1^v^1^;P[pCFD5 dib.KO dgRNA]attP40* (dU6-IRP1A^gR13D^) were generated by our lab. *spok_DmC/TM3,Ser.GFP* (spok_DmC), *y^1^v^1^;P[pCFD5 dib.KO dgRNA]attP40* (dU6-dib^gR1^), *y^1^v^1^;P[pCFD5 dib.KO dgRNA]attP40* (dib TSS^−110^), *y^1^v^1^;P[pCFD5 dib.KO dgRNA]attP40* (dU6-phm^gR1^), *P[pCFD5 dib.KO dgRNA]attP40* (dU6-IRP1A^gR^)*, IRP1A^KO^/TM6B, Hu, Tb* were previously generated by our lab [8, 74]. *y^1^w*P[nos-PhiC31.NLS]X; attP40(II)* and *y^1^w*P[nos-PhiC31/int.NLS]X; attP2(III)* were gifts from the BestGene Inc. *phm22-Gal4* was a kind gift from Michael O’Connor’s lab. Stocks were maintained on a cornmeal diet unless otherwise specified.

### Survival studies

Experiments were carried out as previously described [8,59,74]. In brief, 50 embryos per replicate were collected in 1-hour intervals and transferred to vials containing appropriate media. Larval survival was scored for every stage. At least three independent crosses (= three biological replicates) were carried out per experimental condition. Modified media were prepared by adding compounds (e.g., iron or 20E) during the preparation process. For iron-enriched media, a 1 M stock solution of Ferric Ammonium Citrate (FAC) (Sigma #F5879) was used to make a medium with a final concentration of 1mM FAC. For 20-Hydroxyecdysone (20E)-supplemented media, the final concentration was 0.33 mg/ml. For *dib^2^*, *phm^E7^* mutants or transgenic lines that ubiquitously knock-down *dib* or *phm*, fresh embryos were immersed for 5 min in 1xPBS containing 20E at the final concentration of 0.11 mg/ml [59]. Survival rates were normalized to the number of embryos used per replicate. Error bars represent standard deviation (data is normally distributed).

### Embryo injection

PhiC31 constructs were injected at 500-600 ng/μl concentrations. Injections were performed either at the University of Alberta or Da Lat University using standard procedures [115]. 300-500 embryos were injected per construct. Surviving adults were backcrossed to *w^1118^* (for Cas13 transgenes) or *y^2^cho^2^v^1^* (for crRNA transgenes) and used to generate independent lines.

### Quantitative real-time PCR (qPCR)

Studies were performed as previously described [8, 74]. The extracted RNA (Qiagen RNeasy extraction kit) was reverse-transcribed via the ABI High Capacity cDNA synthesis kit (ThermoFisher #4368814). Synthesized cDNA was used for qPCR (QuantStudio 6 Flex) using KAPA SYBR Fast qPCR master mix #Sigma KK4601). For each condition, three biological samples were each tested in triplicate. Samples were normalized to *rp49* based on the ΔΔCT method. Error bars represent 95% confidence intervals.

### Statistical analysis

For the survival studies, survival rates were normalized to the starting number of embryos (50 embryos per replicate). Error bars represent standard deviation (data is normally distributed). In the FREPAIRv2 editing experiment, the editing rate was calculated as the percentage of samples with correct modification out of the total number of sequenced samples. The off-target rate represents the percentage of samples with incorrect modifications out of the total number of sequenced samples. Error bars represent standard deviation (data is normally distributed). In qPCR reactions, samples were normalized to *rp49*, a housekeeping gene, and based on the ΔΔCT method [116], error bars represent 95% confidence intervals and contain the error for the calibrator (which is shown without error bars). For multiple comparisons to the same control, we used one-way ANOVA, followed by Dunnett’s test. For multiple pair-wise comparisons (in the RNA immunoprecipitation experiments), we applied one-way ANOVA coupled with Tukey’s HSD (HSD = honestly significant difference) test. At least three biological samples and three technical replicates were analysed per condition. For quantification of the eCFP signal in immunostains or western blots, the mean pixel values of the images were analyzed using ImageJ as the corrected total cell fluorescence (CTCF) using the formula: CTCF = selected cell intensity – (area of the chosen cell * background intensity). The CTCF values were averaged from all biological replicates and normalized to the average CTCF values of no-targeting (NT) crRNA samples. Graphs, standard error calculations, t-tests, Dunnett’s tests, and Tukey HSD were conducted in Microsoft Excel, SPSS (IBM), and Prism 8 (GraphPad). All data were normally distributed.

**Abbreviations**
20E: 20-hydroxyecdysone
ADAR2: Adenosine Deaminase Acting on RNA 2
Cas: CRISPR-associated proteins
CasRX: *Ruminococcus flavefaciens* XPD3002 Cas13d
CoIP: coimmunoprecipitation
CRISPR: cluster regularly interspaced short palindromic repeats
crRNA: CRISPR RNA
dCas9: nuclease-dead Cas9
*dib*: *disembodied*
dU6:3: *Drosophila* U6:3 promoter
eCFP: enhanced cyan fluorescent protein
ENC: effective number of codons
gRNA: guide RNA
IRE: iron-responsive element
IRP: iron regulatory protein
L1: first instar larvae
L2: second instar larvae
L3: third instar larvae
LshCas13a: *Leptotrichia shashii* Cas13a
LwaCas13a: *Leptotrichia wadei* Cas13a
*mt:CoI*: *mitochondrial cytochrome c oxidase subunit I*
*mt:CoII*: *mitochondrial cytochrome c oxidase subunit II*
mtDNA: mitochondrial DNA
NES: nuclear export signal
NLS: nuclear localization signal
NT: non-targeting
PspCas13b: *Prevotella buccae* Cas13b
RNAi: RNA interference
SHERLOCK: specific high-sensitivity enzymatic reporter unlocking

## Declaration

### Ethics approval and consent to participate

Experiments in this study were conducted in *Drosophila* cells and live organisms following standard protocols. No human samples were used. No ethics approval is required in this study.

### Consent for publication

All authors participated in this study have been notified about the preparation and submission of the manuscript. Data generated by each author have been collected and used for the preparation of this manuscript.

### Availability of data and materials

The datasets supporting the conclusions of this article are available in the Source Data file at Figshare repository (https://figshare.com/s/ec49a9766bffb1bfe712) with DOI information https://doi.org/10.6084/m9.figshare.12702317. The pC13cr01 vector collection for cell culture work, p13X and p13B vectors for generating *in vivo* crRNA transgenes have been deposited at the *Drosophila* Genetic Resource Center (DGRC) and tentatively scheduled to be available in December 2020. Fly strains carrying CasFB and CasFX will be available at the Bloomington *Drosophila* Stock Center starting January 2021 as a part of the National Institute of Health project (NIH P40ODO18537). Import permit for cell lines Sg4_CD and Sg4* has been obtained and the live stocks will be sent to DGRC in October 2020. Fly strains, plasmids, and cell lines can also be obtained from our lab upon request.

## Author Contributions

N.H. co-designed & carried out most of the experiments, and wrote the manuscript. N.D. generated parts of the pC13cr01 plasmids used for cell culture. R.L. assisted with the generation of Cas13 transgenes. K.K.J. acquired funding, supervised trainees, co-designed experiments, and revised the manuscript.

## Acknowledgments

We thank Norbert Perrimon, Simon Bullock, Gerald Rubin, Feng Zhang, Christoph Metzendorf, Kate O’Connor, James D. Sutherland, Andrew Simmonds, and Patrick Hsu for sharing the original plasmids used in this study. Some stocks used in this study were obtained from the Bloomington *Drosophila* Stock Center (NIH P40ODO18537), the Vienna *Drosophila* Resource Center, the Japan National Institute of Genetics. This work was supported by the Natural Sciences and Engineering Research Council of Canada (NSERC #RGPIN-2018-04357) and the Canadian Institute for Health Research (CIHR #PS 169196).

---

**Figure S1:**
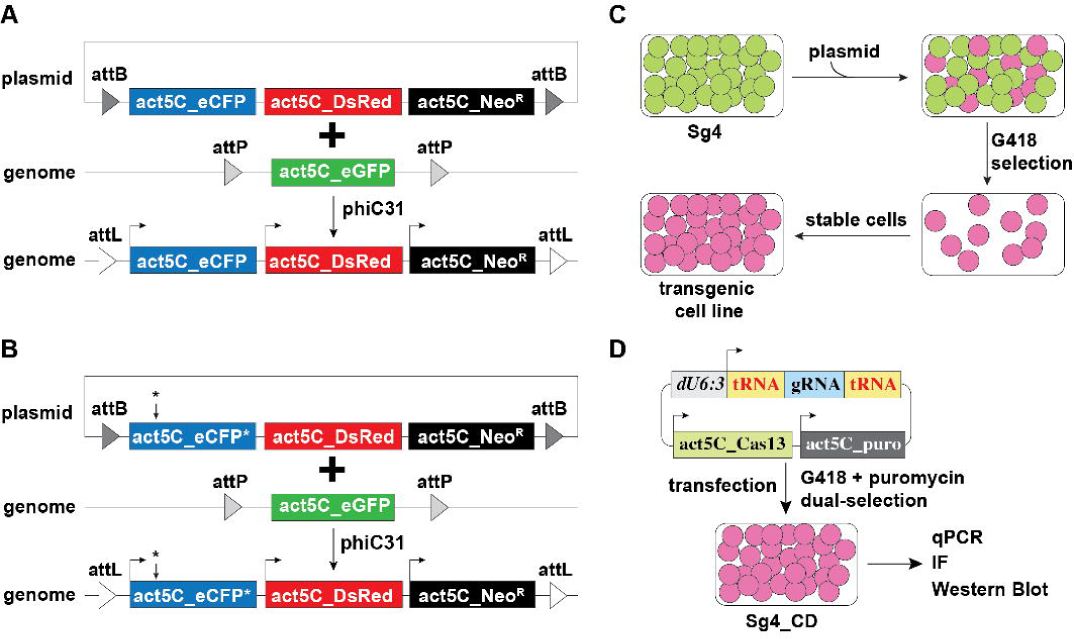
Schematic of transgenic cell culture and *in vitro* study. **(A)** Generation of Sg4_CD cell line that expresses *eCFP*, *DsRed*, and *Neo^R^* genes under independent *actin5C* (*ac5*) promoters. **(B)** Generation of Sg4* cell line that expresses mutant *eCFP\**, *DsRed*, and *Neo^R^* genes under independent *actin5C* (*ac5*) promoters. **(C)** Establishment of the transgenic cell line. Two days after transfection, cells were supplemented with geneticin (G418). Cells without transfected plasmid will be eliminated eventually, leaving cells with successful integration. Cells were passaged for at least four rounds, and integration was confirmed via sequencing. **(D)** Schematic of pC13cr01 vector activity. Upon transfection with the pC13cr01 vector, the cells were selected with geneticin and puromycin to eliminate untransfected cells. Seven days after transfection, cells were collected for later study.

**Figure S2:**
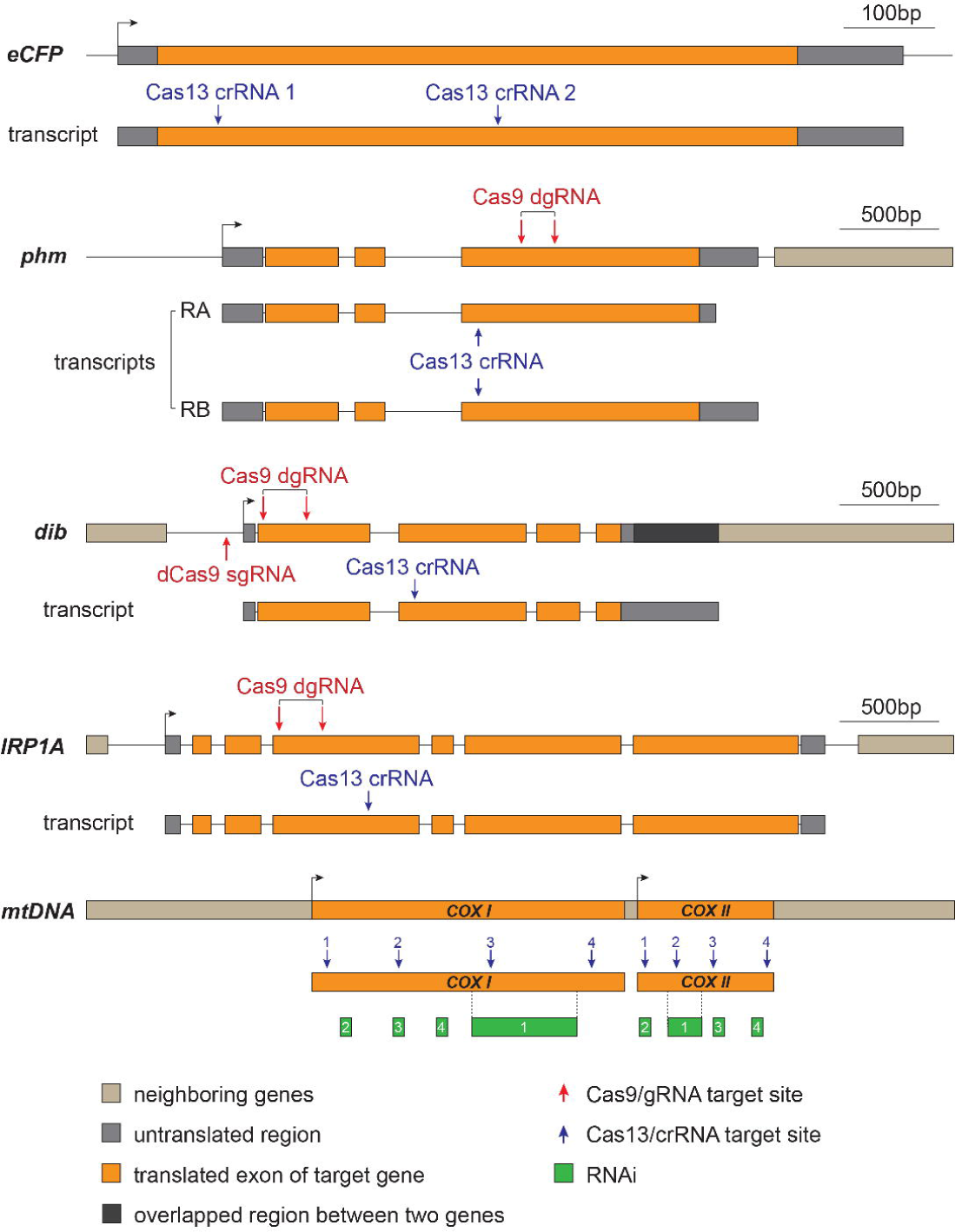
Target sites of crRNAs. For *in vitro* evaluation, we tested *eCFP* expression and two mitochondrial-encoded transcripts, *COXI* and *COXII*. For the *in vivo* approach, we tested two genes that encode enzymes acting as ecdysteroid-synthesizing enzymes in the *Drosophila* prothoracic gland, *phantom* (*phm*) and *disembodied* (*dib*) and a gene involved in cellular iron homeostasis, namely *iron regulatory protein 1A* (*IRP1A)*. Shown here are the target sites for crRNA (Cas13-compatible, blue), gRNA (Cas9-compatible, red), and RNAi (green) for transcripts we tested.

**Figure S3:**
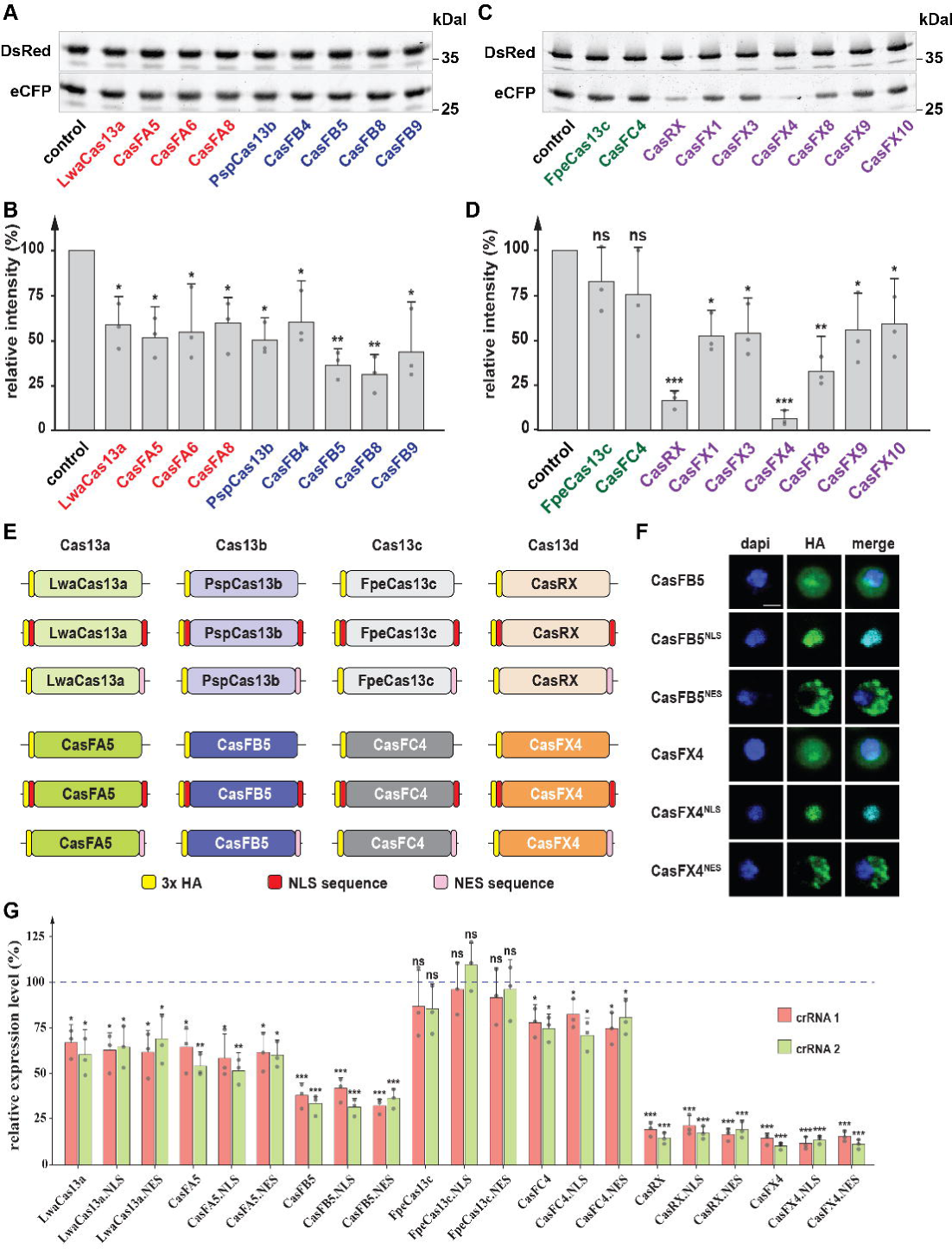
Evaluation of *Drosophila* codon-optimized Cas13 variants. **(A-D)** Western blotting of eCFP in samples that were treated with Cas13 variants that showed the highest efficiency in qPCR experiments. Band intensities were quantified with ImageJ and normalized to samples treated with blank crRNA. * = p-value < 0.05, ** = p-value < 0.01, *** = p – value < 0.001, p-values based on Dunnett’s *post-hoc* test, error bars represent standard error. **(E)** Schematic of Cas13 variants with different signaling sequences, including the <u>n</u>uclear localization <u>s</u>ignal (NLS) and the <u>n</u>uclear <u>e</u>xport <u>s</u>ignal (NES). **(F)** Subcellular localization of CasFB5 and CasFX4 variants in the presence or absence of NLS and NES. Nuclei were stained with dapi (blue) and Cas13 polypeptides were stained with anti-HA antibody (green). Scale bar = 50 μm. **(G)** Evaluation of NLS and NES on Cas13 efficiency via qPCR. Each Cas13 variant was either fused with an NLS or NES and tested for their interference efficiency on eCFP expression. Data were normalized to samples treated with blank crRNA (blue dotted line = 1). * = p-value < 0.05, ** = p-value < 0.01, *** = p-value < 0.001, ns = not significant, p-values based on Student t-tests, error bars represent 95% confidence intervals.

**Figure S4:**
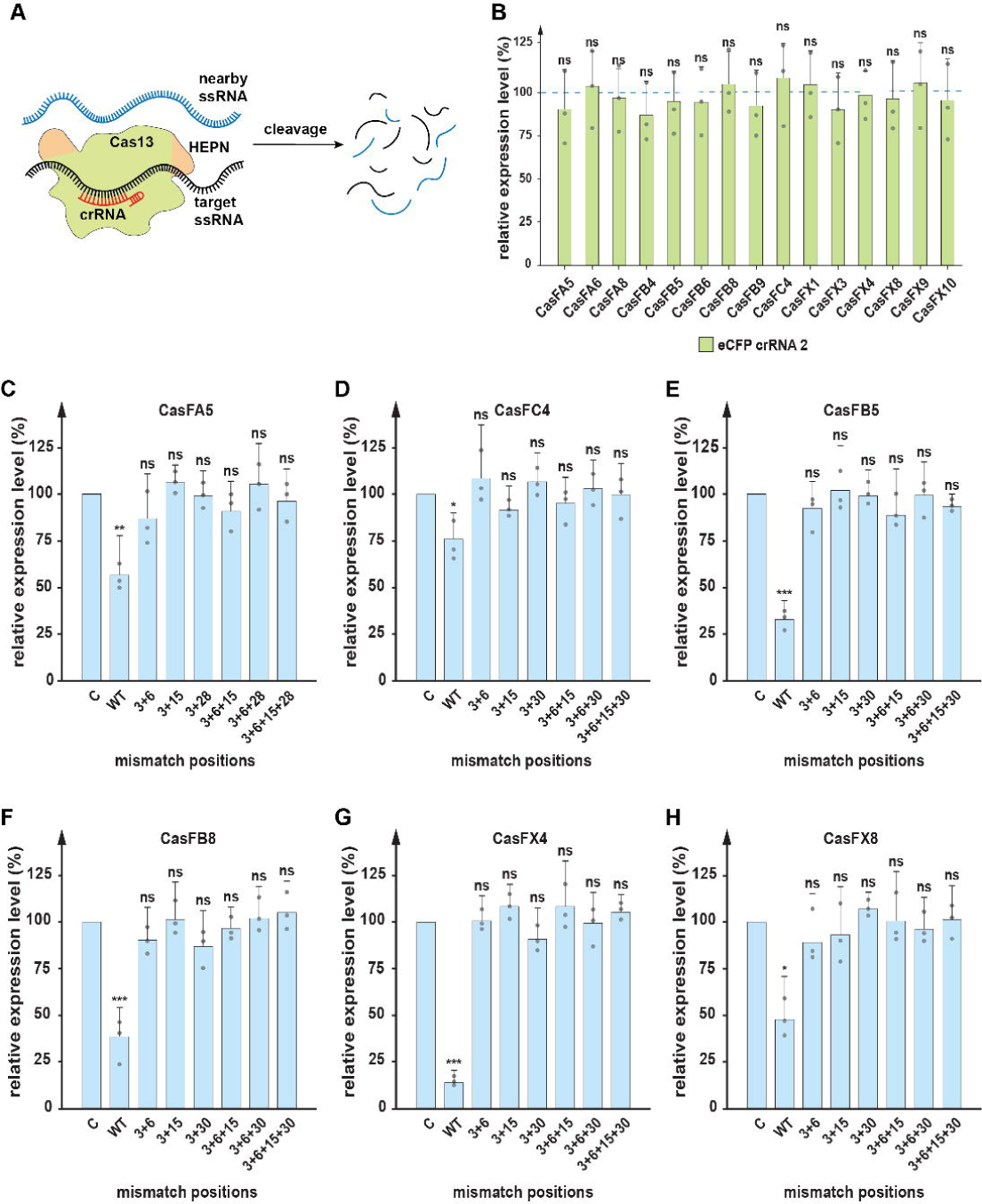
Collateral activity and specificity evaluation of Cas13 variants. **(A)** Schematic of collateral activity in Cas13. Overall, once a complex is formed with its crRNA, and upon binding to target transcripts, Cas13 will undergo a conformational change, which results in the exposure of two nuclease domains (HEPN). This exposure allows the domains to interact with nearby non-specific transcripts and results in their degradation. **(B)** Relative expression of *DsRed* in samples treated with Cas13/crRNA against eCFP. The *act5*C promoter drives DsRed. It is believed that the DsRed transcript is present in high amounts, and is more likely to interact with Cas13. Therefore, if the collateral activity is an issue, we would be expected that DsRed transcript levels are affected. Data were normalized to samples treated with blank crRNA (blue dotted line = 1). * = p-value < 0.05, ** = p-value < 0.01, *** = p-value < 0.001, ns = not significant, error bars represent 95% confidence intervals. **(C-H)** Relative expression levels of eCFP that were exposed to different Cas13 variants and crRNAs carrying different combinations of mismatches along the eCFP crRNA 2. Data were normalized to samples treated with blank crRNA (control = C). eCFP expression level in Cas13/wild-type (WT) crRNA samples were also included as a reference for changes. * = p-value < 0.05, ** = p-value < 0.01, *** = p-value < 0.001, ns = not significant, p-values based on Dunnett’s *post-hoc* test, error bars represent 95% confidence intervals.

**Figure S5:**
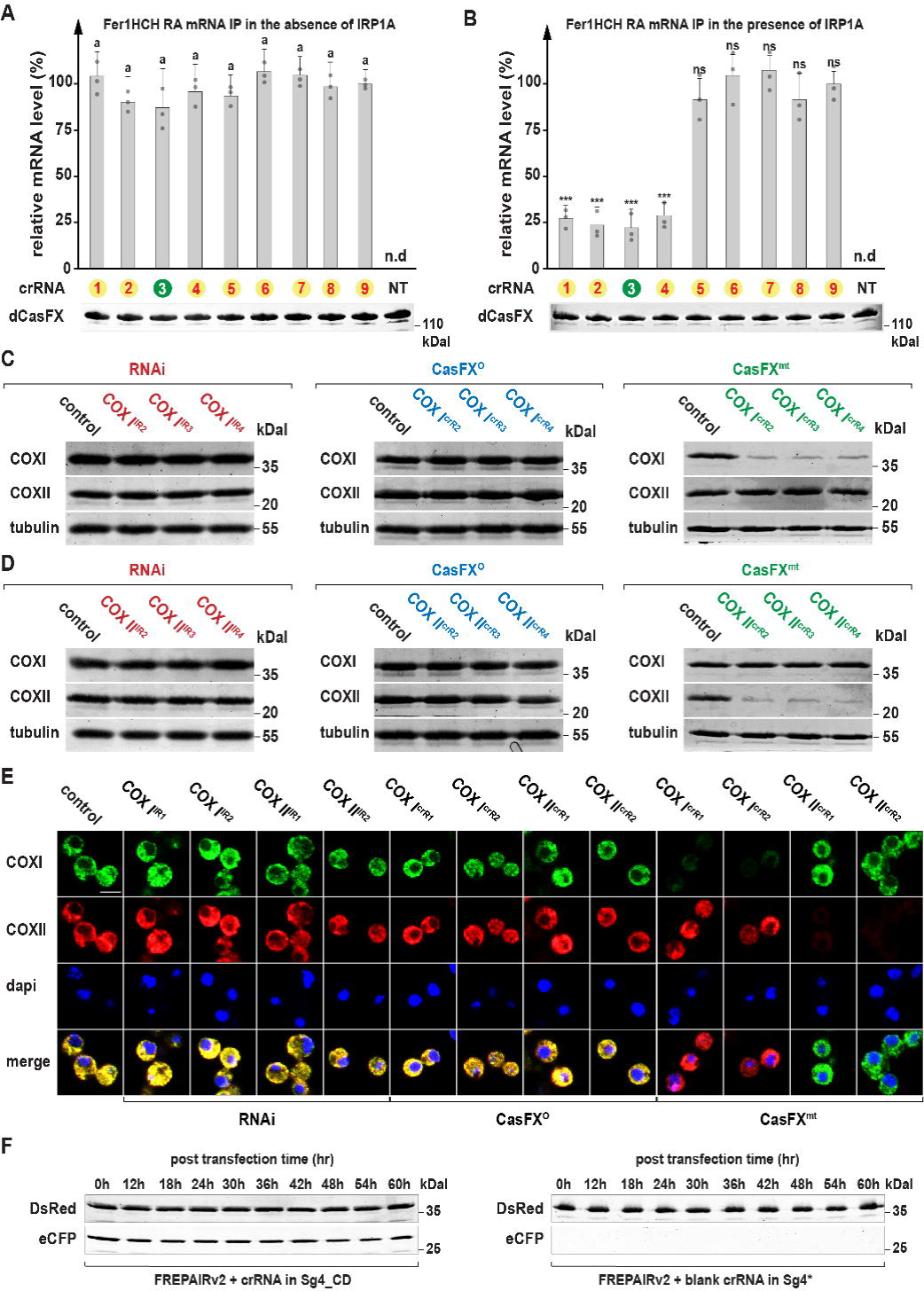
Evaluation of modified CasFX for different approaches. **(A-B)** Relative *Fer1HCH-RA* mRNA amount that was pulled down by the dCasFX/crRNA complex. **(A)** dCasFX and the crRNA targeting *Fer1HCH-RA* mRNA (Figure 4E) were transfected together into one sample, while *Fer1HCH-RA* was transfected into a different sample of cells. The two samples were lysed and combined, followed by immunoprecipitation (IP) of dCasFX via its added HA tag to test for the presence of *Fer1HCH-RA* mRNA. Results were analyzed by one-way analysis of variance (ANOVA) followed Tukey HSD (HSD = honestly significant difference) *post-hoc* test: groups with different letters are statistically different (*p* ≤ 0.05) and groups with the same letters are statistically equal (*p* ≤ 0.05), error bars represent 95% confidence intervals. (**B)** dCasFX and crRNA targeting *Fer1HCH-RA* mRNA (Figure 4E) were transfected together into one sample. *Fer1HCH-RA* and IRP1A^C450S^, the constitutively RNA-binding form of IRP1A that interacts with the iron-responsive element (IRE) in the *Fer1HCH-RA* mRNA, were transfected together into a separate batch of cells. The two samples were lysed and combined, followed by immunoprecipitation (IP) of dCasFX via its attached HA tag to test for the presence of IRP1A in the pull-down assay (Figure 4E) and *Fer1HCH-RA* transcript. Results were analyzed by one-way analysis of variance (ANOVA) followed Tukey HSD (HSD = honestly significant difference) *post-hoc* test: groups with different letters are statistically different (*p* ≤ 0.05) and groups with the same letters are statistically equal (*p* ≤ 0.05), error bars represent 95% confidence intervals. **(C-D)** Western blotting of COXI and COXII targeted by either RNAi, CasFX^O^, or CasFX^mt^. **(E)** Immunofluorescence of COXI and COXII targeted by two independent RNAi constructs, CasFX^O^ or CasFX^mt^. Nuclei were stained with DAPI (blue), COXI was stained with anti-COXI antibody (green), and COXII was stained with anti-COXII antibody (red). Scale bar = 50 μm. (**F)** Western blotting of wild-type eCFP or mutant eCFP* with blank crRNA under the same condition as FREPAIRv2.

**Figure S6:**
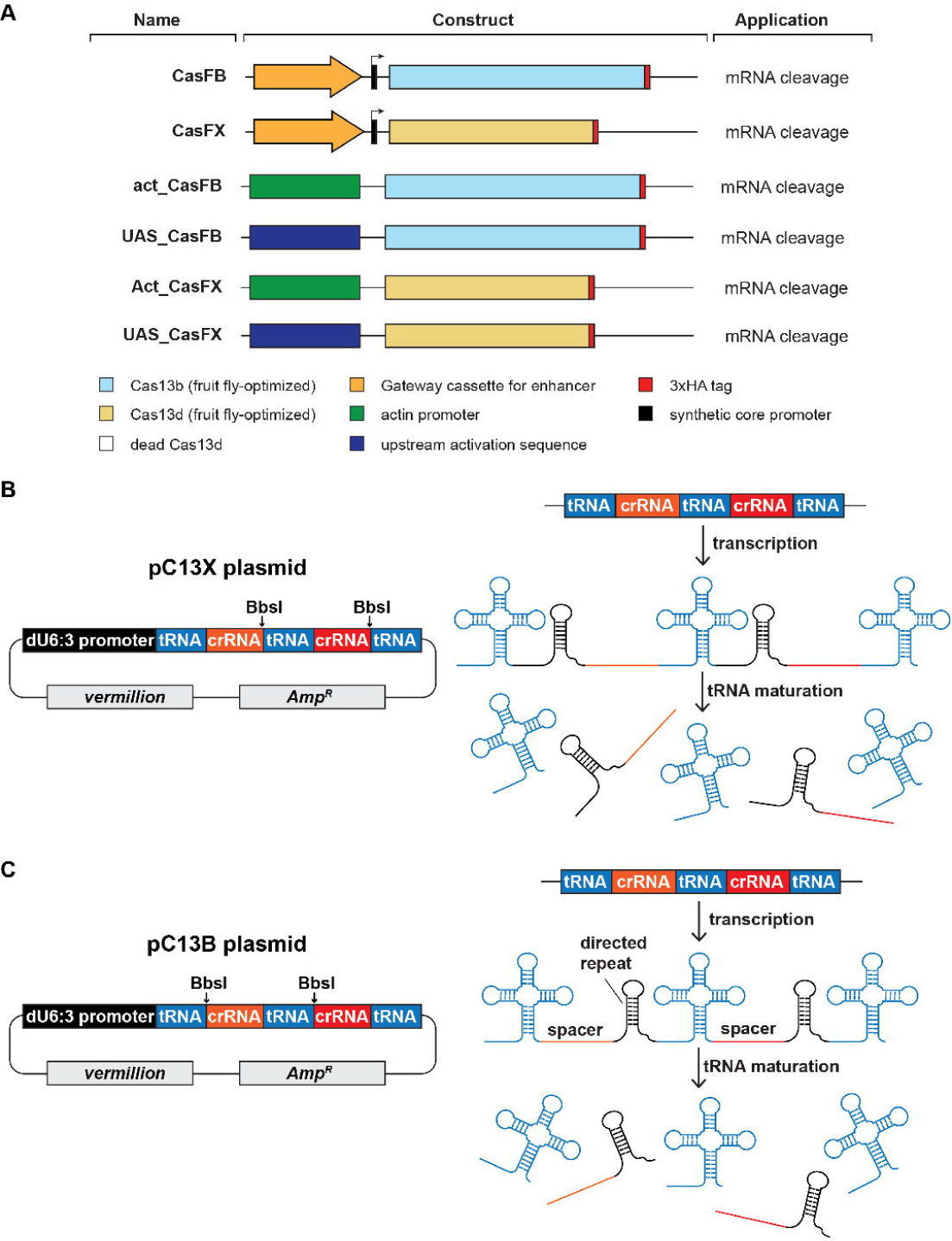
CRISPR/Cas13 transgenes and crRNA vectors for *in vivo* RNA targeting. **(A)** Collection of Cas13 transgenes. The general Cas13 collection is composed of a *mini-white* gene as a marker, a PhiC31 integrase-compatible *attB* site, and the *bla* coding sequence to mediate ampicillin resistance and a synthetic core promoter. Shown here are the gateway cassette for an enhancer of choice, and the Cas13 variants. The gateway cassette allows using LR Clonase-based recombination (ThermoFisher) to insert enhancer/promoter regions to drive tissue-specific *Cas9* expression. The *act-Cas13* transgenes drive the expression of *Cas13* via *actin 5C* (*ac5*) promoter while the *UAS-Cas13* transgenes allow tissue-specific expression of *Cas13* via the Gal4/UAS system. In all cases, Cas13 variants were fused with a 3xHA epitope tag at the C-terminal end. **(B)** Collection of Cas13-compatible crRNA vectors. pC13X is compatible with CasF, whereas pC13B is designed for CasFB. Both vectors carry a *vermillion* marker, a PhiC31 integrase-compatible *attB* site, and the *bla* coding sequence to mediate ampicillin resistance. Each vector holds a multiplex tRNA:crRNA cassette to facilitate the cloning of corresponding crRNA via BbsI digestion. The cassette is driven by the ubiquitous *Drosophila* U6:3 promoter (dU6:3) and is transcribed as a single transcript. Upon tRNA maturation, crRNA will be released and ready to form a complex with Cas13 nuclease.

**Figure S7:**
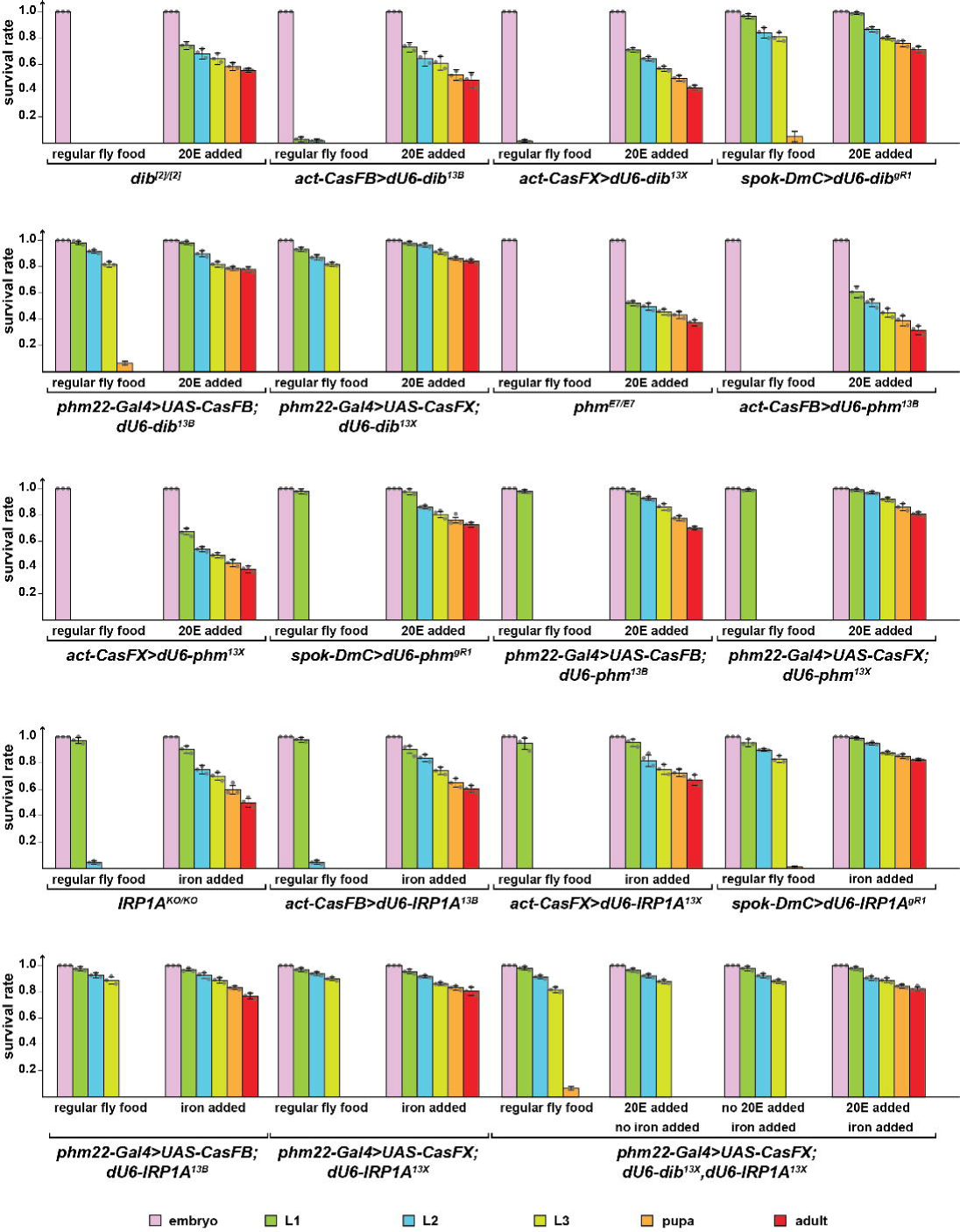
*In vivo* efficiency of *Drosophila* codon-optimized CRISPR/Cas13 variants. Survival rates of classic mutants (*disembodied: dib^2^, phantom: phm^E7^*, *IRP1A*: *IRP1A^KO^*), reared on either regular fly food, fly food supplemented with 20-Hydroxyecdysone (20E), or fly food supplemented with iron. Expression of transgenes was either driven by Gal4 (*phm22-Gal4* for prothoracic gland-specific expression) or by direct regulation by an enhancer (*act-CasFX*; *act-CasFB* for ubiquitous expression and *spok-DmC* for prothoracic gland-specific expression). CasFX and CasFB are Cas13 variants from this study, while *spok-DmC* drives the expression of CRISPR/Cas9 [8]. *dU6-dib^gR1^*, *dU6-phm^gR1^* and *dU6-IRP1A^gR1^* are ubiquitously expressed sgRNAs used for CRISPR/Cas9-mediated gene disruption, while all other *dU6* transgenes express crRNAs that work in conjunction with Cas13. Data were normalized to the number of embryos in the starting population. Error bars represent standard deviation.

**Figure S8:**
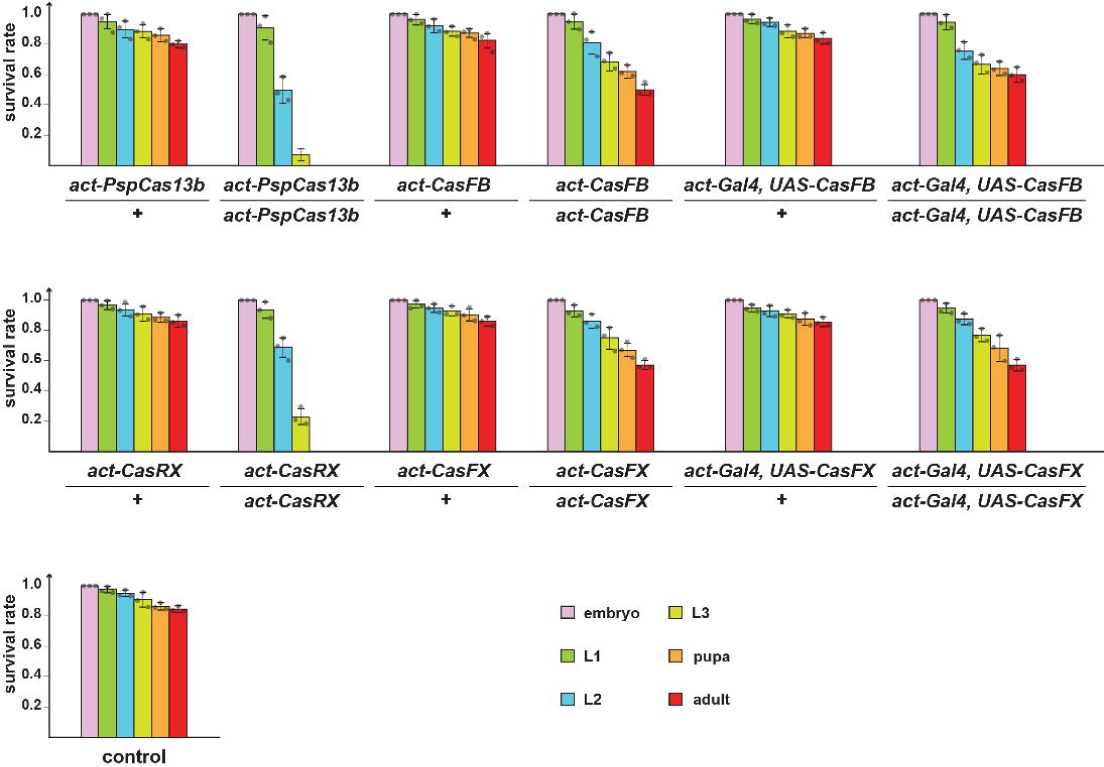
Survival rates of transgenic *Drosophila* lines carrying codon-optimized CRISPR/Cas13. Survival rates of populations heterozygous or homozygous for Cas13 transgenes, including *act-PspCas13b*, *act-CasFB*, *act-Gal4*>*UAS-CasFB*, *act-CasRX*, *act-CasFX*, and *act-Gal4*>*UAS-CasFX*. Survival rates of the *w^1118^* strain were used as a control. Data were normalized to the number of embryos used in the starting population. Error bars represent standard deviation.

**Table S1: *Drosophila* codon-optimized Cas13 variants.**

**Table S2: List of plasmids.**

**Table S3: Survival data of all Cas13/crRNAs lines.**

**Table S4: Primers.**

**Supplemental methods S1: Cloning procedures for CRISPR/Cas13-crRNA vectors.**

